# A tale of too many trees: a conundrum for phylogenetic regression

**DOI:** 10.1101/2024.02.16.580530

**Authors:** Richard Adams, Jenniffer Roa Lozano, Mataya Duncan, Jack Green, Raquel Assis, Michael DeGiorgio

## Abstract

Just exactly which tree(s) should we assume when testing evolutionary hypotheses? This question has plagued comparative biologists for decades. Given a perfectly estimated tree (if this is even possible in practice), we seldom know with certainty whether such a tree is truly best (or even adequate) to represent the evolutionary history of our studied traits. Regardless of our certainty, choosing a tree is required for all phylogenetic comparative methods. Yet, phylogenetic conflict and error are ubiquitous in modern comparative biology, and we are still learning about their dangers when testing evolutionary hypotheses. Here we investigated the consequences of gene tree-species tree mismatch for phylogenetic regression in the presence of incomplete lineage sorting. Our simulation experiments reveal excessively high false positive rates for mismatched phylogenetic regression with both small and large trees, simple and complex traits, and known and estimated phylogenies. In some cases, we find evidence of a directionality of error: incorrectly assuming a species tree for traits that evolved according to a gene tree sometimes fares worse than the opposite. To explore difficult yet realistic regression scenarios, we also used estimated rather than known trees to conduct case studies, as well as an expansive gene expression dataset to investigate an arguably best-case scenario in which one may have a better chance to match tree with trait. Though never meant to be a panacea for all that may ail phylogenetic comparative methods, we found promise in the application of a robust estimator as a potential, albeit imperfect, solution to some issues raised by tree mismatch, perhaps offering a path forward. Collectively, our results emphasize the importance of careful study design for comparative methods, highlighting the need to fully appreciate the role of adequate phylogenetic modeling for testing evolutionary hypotheses.

## Introduction

It is a tale nearly as old as time: you measure a set of traits across a sampling of organisms, and you seek to gain some new biological insights by testing for statistical relationships between two or more of your studied traits. Both scientists and philosophers alike have strived to understand biodiversity for centuries now by employing such a strategy. The diversity of traits that can be studied span any characteristic that can be reliability measured from organisms, from individual cells (e.g., cell size, cell morphology, and gene expression; Chen et al. 2023; Gu 2016; Dunn et al. 2018) to organismal-level traits (e.g., body size, head morphology, and behavior; Al-Kahtani et al. 2004; Ross et al. 2004; Kamilar and Cooper 2013); we are typically only limited by our own curiosity, time, and funding perhaps. For this study, we focus on quantitative traits of varying architectures. For example, does body size predict brain size? Does the expression of one gene predict the expression of another? Does propagule size predict invasiveness? The possibilities seem almost endless, and many classical statistical approaches to linear regression appear well-suited to these questions. Because we are living in the 21st century, we also know that phylogeny must be addressed if we seek reasonable and rigorous answers (Felsenstein 1985; Grafen 1989a; Martins and Hansen 1997; Pagel 1997, 1999a; Rohlf 2001a). What remains less clear, however, is which tree or trees should be considered.

For centuries, comparative studies ignored the role of shared ancestry in shaping observed trends among species and their traits (Felsenstein 1985; Sanford et al. 2002a; Huey et al. 2019). With the advent of phylogenetic comparative methods (PCMs), biologists are now keenly aware of the need to address phylogeny because related species and their traits covary according to their shared ancestry– that is, species are not statistically independent observations (Felsenstein 1985; Grafen 1989a; Martins and Hansen 1997; Pagel 1997, 1999a; Rohlf 2001a). If ignored, then among-species covariance can lead us far astray (Felsenstein 1985; Maddison and FitzJohn 2015a; Uyeda et al. 2018a; Gardner and Organ 2021). By providing a mathematical defense against shared ancestry, phylogenetic regression has become an icon of the comparative method, and its principles have been debated, refined, supplemented, and expanded to target diverse questions, hypotheses, and data types over the years (Harvey and Pagel 1991a; Sanford et al. 2002b; Blomberg et al. 2003a; Felenstein 2004; O’Meara et al. 2006; Revell et al. 2008; Beaulieu et al. 2012; Pennell and Harmon 2013; Maddison and FitzJohn 2015b; Uyeda et al. 2018b). Few studies in evolutionary biology are now published without at least a reference to PCMs and their foundations in phylogenetic regression.

Of course, a fundamental assumption of phylogenetic regression and PCMs generally is that the required input tree is known completely. For real trait data sampled from nature, this assumption is often difficult if not impossible to confirm (Schluter 1995). Simple phylogenetic estimation error is likely to be an issue; errors tend to beget errors, such that errors in the assumed topology and branch lengths may propagate errors in downstream evolutionary inferences that assume error-free trees (Diaz-Uriarte and Garland Jr 1996, 1998; Symonds 2002a; Stone 2011; Mendes et al. 2018). Thus, if an assumed phylogeny is unreliable, then inferences of trait evolution based on Brownian motion and its extensions (Lande 1979; Hansen 1997; Pagel 1999b; Blomberg et al. 2003b; Harmon et al. 2010) may also be suspect (Harvey and Pagel 1991b; Symonds 2002b). Bayesian PCMs that fit models to posterior probability distributions of trees or co-estimate phylogenetic character-evolution parameters hold promise for incorporating estimation uncertainty into the process (e.g., Bastide et al. 2020; Zhang et al. 2021).

Yet, mismatch between an assumed and true phylogeny can occur for reasons besides just estimation error. Perhaps we are simply looking at the wrong tree. That is, we impose a tree for phylogenetic regression that is completely unrelated to our studied trait, its architecture, or its evolutionary past. Put plainly: how confident are we in our ability to accurately (or at least adequately) match our trait to its true phylogenetic history? Relevant to this question is our understanding of the genetic architecture (the number and identity of loci and their trees) underlying a trait of interest. It has become commonplace to effectively ignore genetic architecture by assuming only one particular tree when fitting phylogenetic regression models; for example, often a single tree (e.g., species tree) is assumed to singularly represent the evolutionary history of a trait. Examples abound in the literature of such studies that impose a single tree for regression of complex quantitative traits with almost assuredly complex architectures, such as body size (Jeschke and Kokko 2009), body length (Adams 2013), body mass (Moyers Arévalo et al. 2020), propagule size (Einum amd Fleming 2007), tooth morphology (Weaver and Wilson 2021), and herbivory (Robinson et al. 2023).

We know that variation in phylogenetic history is ubiquitous, arising naturally from speciation, diversification, and evolution (Maddison 1997; Nichols 2001; Degnan and Rosenberg 2009; Kutschera et al. 2014). Importantly, individual gene trees at specific genetic loci often differ wildly from one another and from the overall species tree as a result of incomplete lineage sorting (ILS; Maddison 1997; Nichols 2001; Degnan and Rosenberg 2009; Hobolth et al. 2011), introgression (Yu et al. 2011; Leaché et al. 2014; Solís-Lemus et al. 2016; Tian and Kubatko 2016; Long and Kubatko 2018), ancestral structure (Slatkin and Pollack 2008; DeGiorgio and Rosenberg 2016; Koch and DeGiorgio 2020), and natural selection (Adams et al. 2018; Borges et al. 2020; He et al. 2020; Wascher and Kubatko 2023). Of these processes, ILS is arguably the most infamous (Maddison 1997; Kubatko and Degnan 2007; Edwards 2009; Liu et al. 2015). One particularly concerning consequence of ILS is hemiplasy (Avise and Robinson 2008), which results from forcing trait data to a mismatched tree that is not the true tree, which can generate false patterns of homoplasy-like evolution (Avise and Robinson 2008) and mislead PCMs (Hahn and Nakhleh 2016b; Mendes and Hahn 2016; Guerrero and Hahn 2018; Mendes et al. 2018, 2019; Hibbins et al. 2020).

In the face of widespread gene tree-species tree conflict, how do we choose a tree or trees for phylogenetic regression? Growing evidence suggests that this decision matters, but it can be difficult to know *a priori* whether to assume a particular gene tree or the overall species tree, a specific set of gene trees, or even every possible gene tree. For example, many studies either assume a single species-level phylogeny that has been estimated using coalescent-based (Doña and Johnson, 2023) or traditional concatenation approaches (Hensen et al. 2023), or a single gene tree (e.g., Ross et al. 2004; Al-Kahtani et al. 2004; Kamilar and Cooper 2013; Adams et al. 2014; Gu 2016; Dunn et al. 2018; Chen et al. 2023) to model trait evolution. However, this assumption may (Dimayacyac et al. 2023) or may not (Hahn and Nakhleh 2016a) be the best strategy. On the other hand, modeling evolution as a function of a particular gene tree may prove beneficial for traits predicted to exhibit a more direct one-to-one correspondence with a single tree, such as expression of a gene largely regulated by *cis* elements near its encoded locus (Chen et al. 2019; Bertram et al. 2022; Bastide et al. 2023; Dimayacyac et al. 2023). Perhaps such scenarios arguably represent a best case in which we might at least hope to match tree with trait. Fitting PCMs to gene expression data has also garnered recent considerable interest for modeling functional genomic evolution across cells, tissues, and species (Chen et al. 2019; Bertram et al. 2022; Adams et al. 2023; Bastide et al. 2023; Dimayacyac et al. 2023). Somewhat surprisingly, a recent study found that modeling gene expression as a function of the overall species tree improved the fit of Ornstein-Uhlenbeck models (Dimayacyac et al. 2023), even when compared to local gene trees. Other traits, however, may be subject to more complex architectures that incorporate multiple genetic loci, each with their own genealogical history. Taking this idea further, some recent models assume that all possible gene trees contribute to a given trait (Mendes et al. 2018; Hibbins et al. 2023), which can help alleviate bias of classical PCMs that assume only a single species tree. Importantly, in these and most related studies, a choice of trees was made.

Perhaps we can depict a spectrum of choices that spans assuming a single tree (e.g., species or gene tree), multiple trees (e.g., multiple gene trees reflecting the additive effect of several loci), and onwards to every possible gene tree. No matter where we believe our trait lies on such a spectrum, we must choose one tree or another (or try multiple options) when applying phylogenetic regression. Thus, we argue that this decision is relevant because practically, PCMs require such a choice, and realistically, because it is seldom possible to know the precise numbers and identities of underlying genetic loci (and their associated trees) encoding the vast majority of traits. For now, let us not get into the frightening possibility that genetic architecture itself may change and evolve along a phylogeny. We will save that for a different study perhaps.

Therein lies a phylogenetic conundrum. We must make a choice of trees for any study that applies phylogenetic regression, and yet, we do not fully appreciate or understand the consequences of an incorrect decision. This study seeks to gauge how much we should be concerned about tree mismatch that is not only possible but arguably probable. We do not attempt to comprehensively explore all possibilities on this spectrum, but instead leverage several case studies involving gene tree-species tree discordance and phylogenetic regression. Specifically, we explore the behavior of phylogenetic regression for testing trait associations when the true and assumed trees are mismatched due to the presence of ILS (Figs. 1 and 2). Whereas PCMs can be relatively robust to certain conditions of tree misspecification (Diaz-Uriarte and Garland Jr 1996, 1998; Symonds 2002; Stone 2011), ILS-induced hemiplasy is a particularly devastating (Mendes and Hahn 2016; Mendes et al. 2016, 2018) and widespread (Copetti et al. 2017) phenomenon. Here we employ a large-scale battery of phylogenetic comparative simulations with varying degrees of mismatch for traits of both simple and complex architectures and with both known and estimated trees. After consideration of our simulations, we explore a best-case scenario for matching tree with trait by using an extensive gene expression dataset sampled across mammals. Collectively, our study sought to illuminate and investigate phylogenetic regression in experimentally challenging yet important scenarios.

**FIGURE 1.**
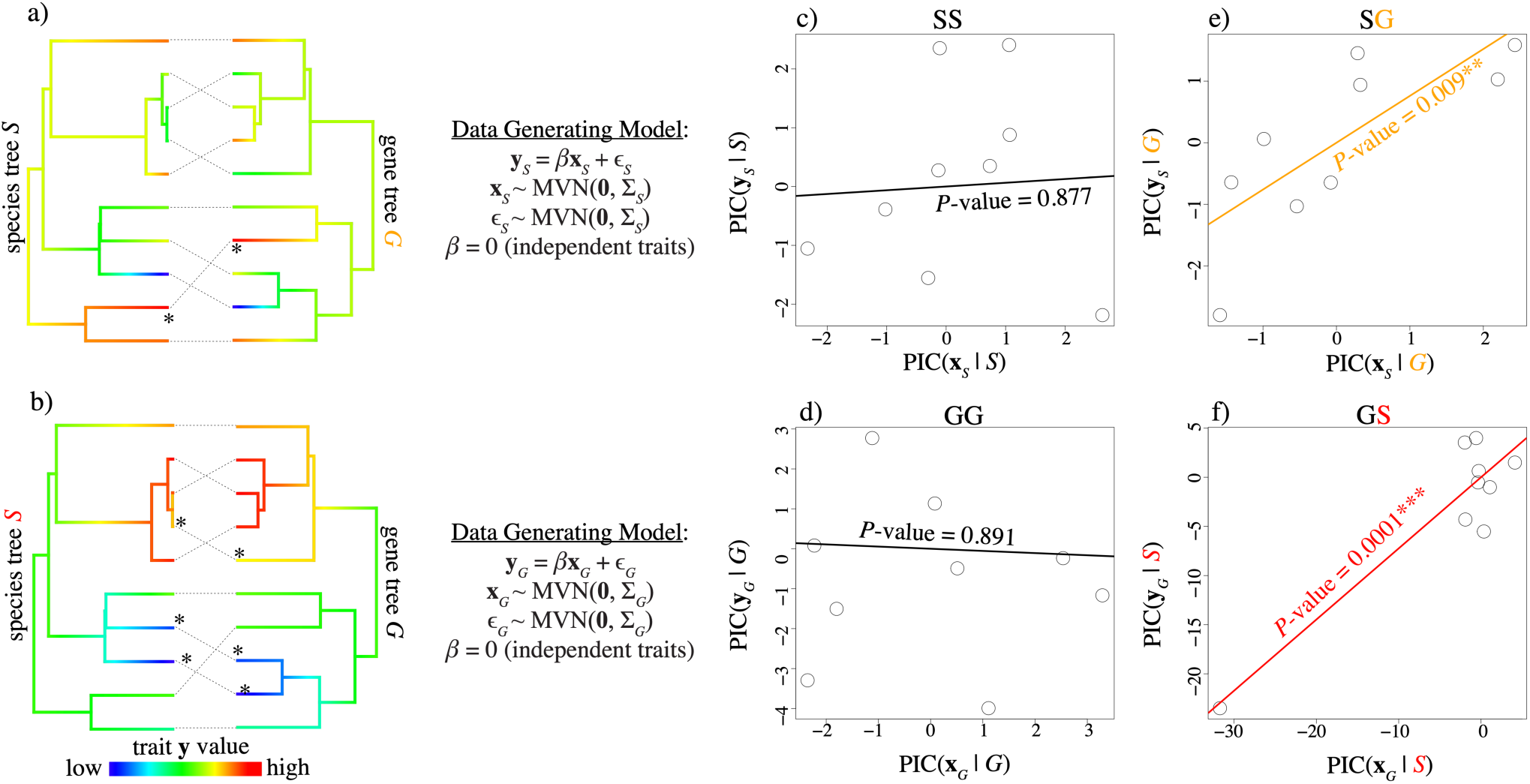
Illustrating the phylogenetic conundrum. Examples showing species tree and gene tree pairs (a and b) and their associated data generating models (center) for scenarios in which traits are generated according to the species tree *S* (top row) or the gene tree *G* (bottom row). Branch colors (a and b) illustrate values of the response trait **y** when mapped to the respective tree using the contMap function from phytools. Two examples (random replicates) of matched phylogenetic regression are shown for SS (c) and GG (d), in which the same tree was used for both generating the trait data and computing PICs, and two examples (random replicates) of mismatched regression are shown for SG (e) and GS (f), in which different trees were used for generating the trait data and computing PICs.

**FIGURE 2.**
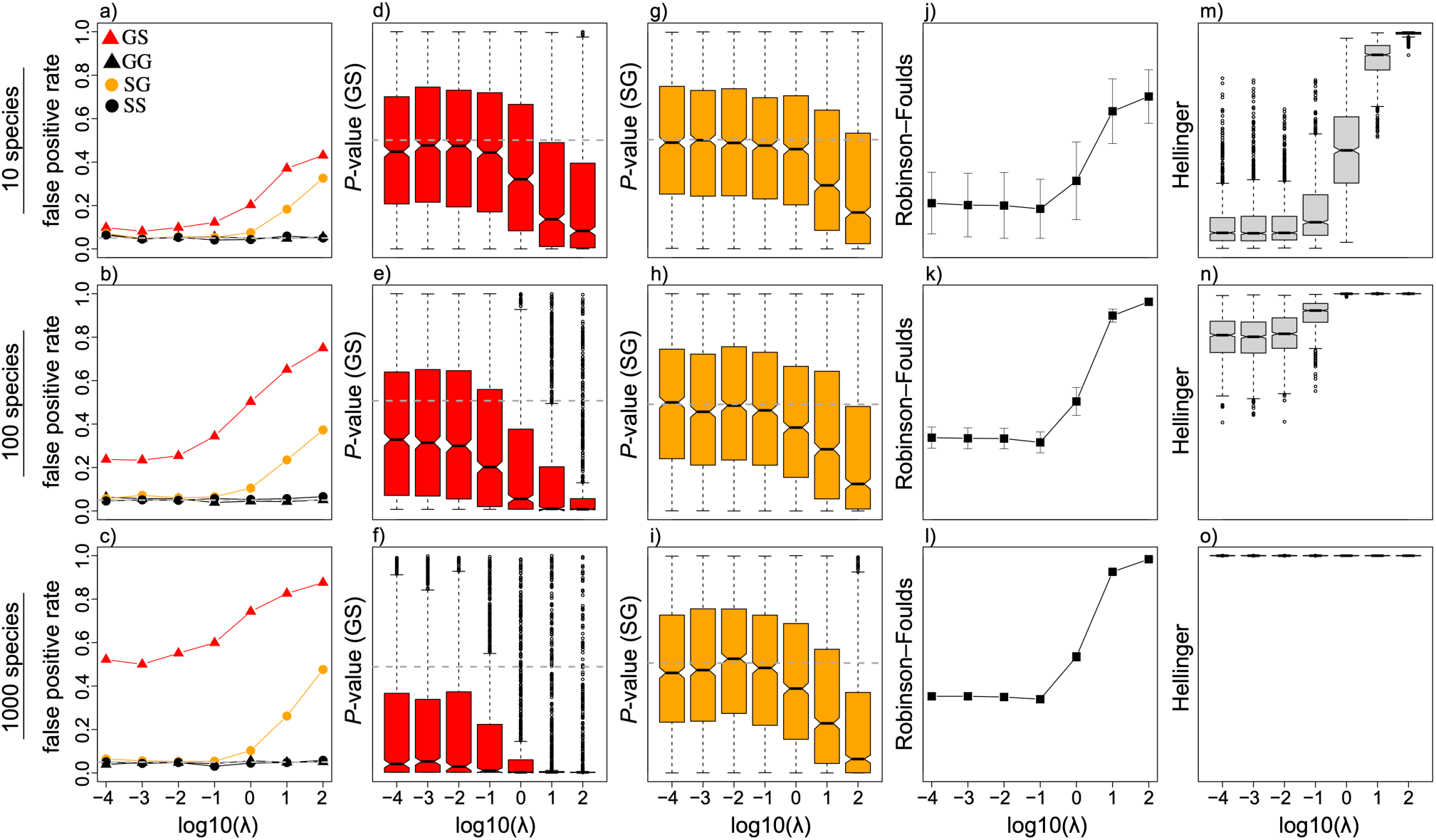
Impacts of mismatched phylogenetic regression. Estimates of the false positive rate (a-c), *P*-value distributions for GS (d-f), *P*-value distributions for SG (g-i), means and standard deviations of Robinson-Foulds topology distances (j-l), and probabilistic Hellinger distances (m-o) between gene trees and species trees from simulations including 10 species (top row), 100 species (middle row), and 1000 species (bottom row) for birth-death simulations with birth rate 𝜆, death rate 𝜆/2, and root age of 10 coalescent units. The two traits were statistically independent (𝛽 = 0) for all simulations. Dashed horizontal lines mark the commonly used false positive rate 𝛼 = 0.05 in panels a-c, median *P*-values taken from matched GG scenarios in panels d-f, and median *P*-values from matched SS scenarios in panels g-i. The *y*-axis ranges from zero to one in all panels.

## Methods

### Simulations with known trees and simple architectures

We explored the performance of phylogenetic regression when using matched versus mismatched trees for testing statistical relationships between two quantitative traits 𝐲 and 𝐱. We generated trait data using a linear model with phylogenetic signal in both the input predictor trait 𝐱 and the response trait 𝐲, following the approach of similar studies of phylogenetic regression (Mazel et al 2016; Revell 2010; Fig. 1). The familiar linear regression equation for two traits can be written as

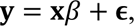

where 𝐲 is an 𝑛-dimensional vector containing measurements of a response trait in each of 𝑛 species, 𝐱 is an 𝑛-dimensional vector containing measurements of an input predictor trait in each of 𝑛 species, 𝛽 is the regression coefficient that measures the relationship between the input predictor trait 𝐱 and the response trait 𝐲, and 𝛜 is an 𝑛-dimensional vector representing the model residuals. Under the null hypothesis of no relationship between 𝐱 and 𝐲, 𝛽 = 0, whereas the alternative hypothesis is that 𝛽 ≠ 0. Ordinary least squares (OLS) assumes that the residuals 𝛜 are independent and identically distributed as normal with mean zero and some estimated standard deviation; this assumption is commonly violated with phylogenetically-structured data, in which traits tend to covary among species. Variants of phylogenetic regression relax this assumption by considering the phylogenetic variance-covariance structure between a set of 𝑛 species that is defined by their evolutionary relationships. Phylogenetic independent contrast computes a set of 𝑛 – 1 contrasts that are statistically independent, whereas phylogenetic generalized least squares directly incorporates the phylogenetic variance-covariance structure in the model. Though differing in their strategies for dealing with phylogeny, 𝛽 estimates and significance levels are equivalent for both approaches under Brownian motion (Blomberg et al. 2012).

To simulate traits according to a tree, we included phylogenetic signal into the linear model by simulating 𝐱 and 𝛜 according to a multivariate normal (MVN) distribution with mean zero and an 𝑛 × 𝑛 phylogenetic variance-covariance matrix 𝚺, which is defined according to a specific species tree or gene tree (Grafen 1989; Martins 1996; Martins and Garland 1991). When denoting the data generating process, we use the subscript *S* for traits with signals matching a species tree *S* and the subscript *G* for traits with signals matching a gene tree *G*. Therefore, to generate trait data with phylogenetic signal according to a species tree *S*, we used

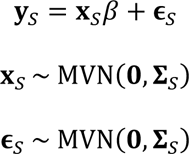

where both 𝐱*_S_* and 𝛜*_S_* are distributed as multivariate normal with mean 𝑛-dimensional vector 𝟎 containing all zero elements and phylogenetic variance-covariance 𝚺*_S_* defined according to the species tree *S*. Note that when 𝛽 = 0, the response trait is simply distributed as 𝐲*_S_* ∼ MVN(𝟎, 𝚺*_S_*), which effectively represents independent Brownian motion evolution for both 𝐲*_S_* and 𝐱*_S_* on the same species tree. Similarly, we generated trait data with phylogenetic signal according to a gene tree *G* using

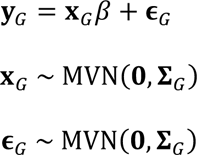

where both 𝐱*_G_* and 𝛜*_G_* are distributed as multivariate normal with mean 𝑛-dimensional vector 𝟎 containing all zero elements and phylogenetic variance-covariance 𝚺*_G_* defined according to the gene tree 𝐺. After generating trait data with these two data generating models, phylogenetic regression was conducted by computing phylogenetic independent contrasts (PICs) using either the species tree *S* or the gene tree *G*, allowing us to explore scenarios of phylogenetic mismatch in which the data generating process and the assumed tree for PICs are different (details provided below). To assess false positive rates under different scenarios, we set the true regression coefficient 𝛽 = 0, whereas a nonzero 𝛽 ≠ 0 was used to investigate statistical power under different scenarios.

To investigate impacts of ILS on phylogenetic regression, we compared “matched” regression (Figs. 1c and d), for which the same tree is used to generate and compute PICs, to “mismatched” regression, for which different trees are assumed to generate and compute PICs (Figs. 1e and f). We conducted a multifactorial simulation study to investigate both the false positive rate and power of phylogenetic regression across a range of scenarios with increasing probabilities of tree mismatch due to ILS. Our overall simulation protocol can be described in four steps: (1) a species tree *S* is generated using a diversification process (Yule 1925a), (2) a gene tree *G* is simulated according to the multispecies coalescent with the species tree *S* obtained from step 1, (3) comparative trait data are simulated using the phylogenetic variance-covariance matrix 𝚺*_S_* from the species tree obtained from step 1 to obtain 𝐲*_S_* and 𝐱*_S_*, *or* using the phylogenetic variance-covariance matrix 𝚺*_G_* from the gene tree from step 2 to obtain to obtain 𝐲*_G_* and 𝐱*_G_*, and (4) phylogenetic regression is conducted with the original algorithm of Felsenstein (1985) on PICs applied to the trait data generated from step 3 (Fig. 1) with either matched or mismatched trees. That is, the opportunity for mismatch occurs in steps 3 and 4 when computing PICs using an incorrect tree that is unrelated to the data generating process of the studied traits. When the same tree is used in steps 3 and 4, the scenario represents matched regression because the tree used to simulate the traits is also used to compute PICs for phylogenetic regression. Conversely, when different trees are used for steps 3 and 4, the scenario represents mismatched regression, as the assumed tree is not the tree that generated the data (e.g., a species tree is assumed for traits simulated on a gene tree).

Our simulation approach therefore allowed us to examine modeling conditions representing four distinct scenarios: matched gene tree-gene tree (GG), matched species tree-species tree (SS), mismatched gene tree-species tree (GS), and mismatched species tree-gene tree (SG), where the first tree in each pair indicates the tree used to simulate traits, and the second tree is assumed when computing PICs for phylogenetic regression (Fig. 1). For example, GG represents the matched scenario for which the same gene tree *G* is used to both generate the two traits and compute PICs for 𝐲*_G_* and 𝐱*_G_*, whereas GS is mismatched because a gene tree *G* is used to generate the traits 𝐲*_G_* and 𝐱*_G_*, but the species tree *S* is incorrectly assumed when computing PICs for regression. Likewise, both the 𝐲*_S_* and 𝐱*_S_* traits and their PICs are generated with the same species tree for SS scenarios, whereas SG represents tree mismatch because the traits 𝐲*_S_* and 𝐱*_S_* are both generated via the species tree, but a gene tree is incorrectly assumed when computing PICs. Thus, we evaluated phylogenetic regression with two forms of correctly specified models (GG and SS) and with two forms of model misspecification with mismatched trees (GS and SG; Fig. 1).

Throughout our simulations, we varied both the total number of taxa 𝑛 ∈ {10, 100, 1000} and speciation rate 𝜆 ∈ {10^-4^, 10^-3^, …, 10^2^} used to simulate the species trees. By varying 𝜆 across this logarithmically scaled range, we effectively incorporated variability in the expected amount of phylogenetic discordance and potential for hemiplasy due to ILS, as 𝜆 is inversely proportional to the expected branch lengths in the species tree. In particular, slow rates (𝜆 = 10^-4^ yield long internal branch lengths and lower ILS, whereas fast rates (𝜆 = 10^2^) generate short internal branch lengths, exacerbating ILS and increasing the probability of gene tree-species tree discordance. We generated species trees under a birth-death model in which the death rate was set to half the speciation rate; we also investigated a simple pure-birth model of diversification (Yule 1925b) with the death rate set to zero. We employed the R package TreeSim (Stadler 2011) using the s*im.bd.taxa.age* function with a most recent common ancestor age of either one, 10, or 100 to generate species trees of varying depths. We used the *sim.coaltree.phylo* function in R package Phybase (Liu and Yu 2010) to simulate gene trees based on species trees. Trait data were then simulated according to either *S* or *G* using the linear models described above for a total of 10^3^ replicates for each value of 𝜆 ∈ {10^-4^, 10^-3^, …, 10^2^} and for each of the four scenarios GG, GS, SS, and SG. For each replicate, phylogenetic regression was conducted using PICs computed according to the four scenarios (Fig. 1) using the *pic* function provided in R package APE (Paradis and Schliep 2019). We evaluated the false positive rates for each scenario by setting the true regression coefficient 𝛽 = 0 and quantifying the number of replicates with *P*-value < 0.05 that incorrectly reject the null hypothesis, whereas four values of nonzero 𝛽 ∈ {0.25, 0.50, 0.75, 1.0} were used to investigate statistical power for correctly rejecting the null hypothesis when 𝛽 ≠ 0. To provide context on the degree of gene tree-species tree discordance, we computed Robinson-Foulds distances (Robinson and Foulds 1981) and probabilistic trait Hellinger distances (Pardo 2005; Adams et al. 2021) between the gene tree and species tree for each replicate. The Robinson-Foulds metric considers only the topological distance between two trees, whereas Hellinger is a probabilistic distance between two MVN distributions, which is applicable to many models of trait evolution based on principles of Brownian motion that model trait distributions as MVN. To confirm our results for species trees generated under a simpler pure-birth model, we also conducted an experiment using the same protocol as above with 𝑛 ∈ {10, 100, 1000}, a pure-birth model, and a tree depth of 10 to assess false positive rates for 10^3^ replicates of each value of the birth rate.

### Simulations with known trees and complex architectures

Our first array of simulations described above apply simple architectures in which traits were generated according to the phylogenetic variance-covariance matrix of only a single species tree or gene tree (Fig. 1). We also conducted a case study that explored more complex architectures in which trait data are generated according to multiple gene trees, which is expected to occur for many quantitative traits. For these simulations, we followed the same general protocol as above with the addition of the seastaR approach (Hibbins et al. 2023), by computing a phylogenetic variance-covariance matrix 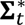 as a weighted mean of the individual gene tree variance-covariance matrices taken from 𝑡 different gene trees. The primary change is that we used 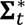 instead of 𝚺*_S_* (a single species tree *S*) or 𝚺*_G_* (a single gene tree *G*) to generate the traits. More specifically, the traits were encoded by 𝑡 gene trees, each with equal weight. We conducted three case study simulations in which the number of gene trees 𝑡 ∈ {2, 5, 10, 100} varied to represent traits with architectures encoded by two, five, 10, or 100 genomic loci and their associated gene trees.

Here we explored matched scenarios in which the same generating 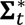 was used to both simulate traits and conduct phylogenetic regression. We also investigated two additional scenarios of mismatched regression: (1) one 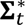 was used to simulate the traits, and a different 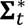 was incorrectly assumed for phylogenetic regression, and (2) one 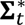 was used to simulate the traits, and the species tree 𝚺*_S_* was incorrectly assumed for phylogenetic regression. We refer to these three scenarios as matched gene trees (i.e., same 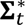 used for both simulation and inference), mismatched gene trees (i.e., different sets of gene trees 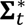 for simulation and inference), and mismatched species tree (i.e., 𝚺*_S_* used for inference instead of the true 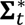), respectively. Because current implementations of PICs require bifurcating trees, we conducted phylogenetic regression using phylogenetic generalized least squares (PGLS; Grafen 1989; Martins 1996; Martins and Garland 1991) using the *gls* function in the R package nlme (Pinheiro et al. 2017). Importantly, the regression slope estimates and levels of significance are equivalent when using phylogenetic regression with PGLS and PIC under Brownian motion (Blomberg et al. 2012). For these analyses, the same birth-death process was used to simulate species tree with a depth of 10 coalescent units and either 10 or 100 species, and we focused on assessing false positive rates when 𝛽 = 0 with 10^3^ replicates for each value of the birth rate 𝜆 ∈ {10^-4^, 10^-3^, …, 10^2^}.

### Simulation case study: How does phylogenetic estimation error influence mismatch?

Results obtained when using true trees may not hold when instead using estimated trees, which is important for empirical studies. Yet simulating species trees, gene trees, and gene sequence alignments entails substantial computational resources. Thus, in addition to simulations that utilized known phylogenies (i.e., those without estimation error), we also conducted a simulation case study that included phylogenetic estimation error for both gene trees and species trees. We followed the above simulation protocol, but included additional steps for estimating gene trees and species trees. Though a multitude of parameters are likely to influence phylogenetic estimation and error, we chose several factors predicted to be important while ensuring computational feasibility. The first three steps of our simulation protocol for this case study are analogous to those of the simulations described above for known trees: (i) simulate a species tree with varying speciation rate 𝜆 ∈ {10^-4^, 10^-3^ …, 10^2^}, (ii) simulate 10 gene trees for each species tree from step i, and (iii) simulate continuous trait using either the known species tree from step i (SS and SG scenarios) or a known gene tree from step ii (GG and GS scenarios). Next, we added components for estimating gene trees and species trees: (iv) simulate 2.5 kilobase alignments for each of the 10 gene trees using an HKY model with a transition/transversion ratio of 4.6 and base equilibrium frequencies of 𝑓_A_ = 0.3, 𝑓_C_ = 0.2, 𝑓_G_ = 0.2, and 𝑓_T_ = 0.3 for nucleotides A, C, G, and T, respectively, (v) estimate gene trees using IQ-TREE (Pei and Wu 2017) with default parameters, (vi) infer a species tree with the gene tree estimates from step v using STELLS2 (Pei and Wu 2017), and (vii) conduct phylogenetic regression using either the estimated species tree from the step vi or the estimated gene tree from step v for computing PICs. In all cases, known phylogenies were used to simulate the trait data (step iii). Therefore, this simulation protocol matches our above simulations with the addition of gene tree estimation (step v) and species tree inference (step vi), with phylogenetic regression conducted using these estimated trees instead of the known trees. For this case study, we focused on the impacts of phylogenetic estimation error on false positive rates of phylogenetic regression by simulating two statistically independent traits with 𝛽 = 0. Because STELLS2 requires at least two samples per species to estimate external branch lengths, the known gene trees simulated in step ii and estimated in step v include two samples per species for species tree estimation. However, for both simulating trait data and fitting regression models, we pruned these trees to include only one lineage sampled per species to allow direct comparisons of phylogenetic regression on gene trees versus species trees with the same numbers of lineages. Because of the computational requirements needed to simulate this multi-step experiment (species tree to gene trees to sequence alignments to inferences of each), we generated 10^2^ replicates for each value of the birth rate parameter 𝜆 ∈ {10^-4^, 10^-3^, …, 10^2^} with 𝑛 = 10 species.

### An empirical best-case study: Does tree choice impact gene expression phylogenetic regression?

Our simulations revealed evidence of profound bias with mismatched phylogenetic regression for traits with both simple and complex architectures, and when using known and estimated trees (see *Results*). Given these findings, we sought to explore the potential empirical impacts of tree choice for a best-case scenario in which one might be able to better match tree with trait. Assuming a single tree (often the species tree) is the typical default for a plurality of studies that employ phylogenetic regression— regardless of trait complexity and whether genetic architecture across a set of species is known or not (e.g., Doña and Johnson, 2023; Hensen et al. 2023; Ross et al. 2004; Al-Kahtani et al. 2004; Kamilar and Cooper 2013; Gu 2016; Dunn et al. 2018; Chen et al. 2023). Thus, we used gene expression as an example of a best-case scenario because one might predict that it should, at least in theory, be easier to match the expression trait value of a given gene to one specific tree—either the species tree or respective gene tree.

We explored the effects of phylogenetic tree specification on tests of trait association using an empirical gene expression dataset from 11 female and male tissues in eight mammals and chicken (Brawand et al. 2011). In particular, we obtained normalized gene expression abundance measurements computed in reads per kilobase of exon model per million mapped reads (RPKM; Mortazavi et al. 2008) from female and male brain (whole brain without cerebellum), female and male cerebellum, female and male heart, female and male kidney, female and male liver, and testis in human (*Homo sapiens*), chimpanzee (*Pan trogodytes*), gorilla (*Gorilla gorilla*), orangutan (*Pongo pygmaeus abelii*), macaque (*Macaca mulatta*), mouse (*Mus musculus*), opossum (*Monodelphis domestica*), platypus (*Ornithorhynchus anatinus*), and chicken (*Gallus gallus*; Brawand et al. 2011). We focused our comparisons by restricting analyses to the most conservative 5,321 orthologous genes, or those with constitutive exons that aligned across all nine species in the original dataset (Brawand et al. 2011), and computed the median expression level for tissues containing multiple replicates. To understand the impacts of tree specification on phylogenetic regression, we obtained the estimated species tree from Brawand et al. (2011) and estimated gene trees from nucleotide and amino acid alignments downloaded from the UCSC Genome Browser (Navarro Gonzalez et al. 2021) at http://www.genome.ucsc.edu. Specifically, the UCSC alignments included all protein-coding exons in human (GRCh38/hg38) and 99 vertebrates (Blanchette et al. 2004; Dreszer et al. 2012), from which we extracted those pertaining to 5,267 genes in the nine species considered here. We concatenated the nucleotide and exon alignments for each gene and constructed gene trees by applying PhyML (Guindon et al. 2010) with default parameters to these alignments. The species tree and all gene trees were scaled to unit depth. We investigated the statistical performance of phylogenetic regression in three experimental settings: expression in female brain ∼ male brain, female heart ∼ male heart, and female kidney ∼ male kidney. For each experiment, we conducted PIC regression based on log-transformed RPKM values across the nine species, and assessed relationships between tissues via evidence of statistical significance (*P*-values).

Specifically, we explored evidence of phylogenetic mismatch when testing for trait covariation between female and male expression across species. For each gene, we conducted phylogenetic regression to test for associations between male and female expression in three separate analyses representing phylogenetic regression based on: (1) the species tree, (2) the gene tree inferred from nucleotide sequences, and (3) the gene tree inferred from amino acid sequences. To explore genome-wide patterns and identify interesting case studies of phylogenetic regression, we also computed three divergence statistics: 𝑑_ST_, 𝑑_NT_, and 𝑑_AT_ representing analyses that assumed the species tree (ST), the nucleotide gene tree (NT), and the amino acid gene tree (AT), respectively. These statistics have the forms

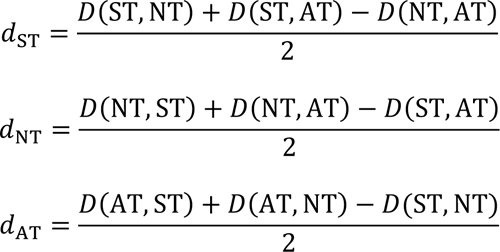

where 𝐷(Tree 1, Tree 2) = |log_10_(𝑃_Tree 1_) − log_10_(𝑃_Tree 2_)| represents the magnitude of the difference between the log-transformed *P*-values of a pair of trees. Thus, each divergence statistic will evaluate whether the *P*-value for a given analysis tree is substantially different from the *P*-values of the other two analysis trees. These measures are akin to those that have been used for identifying population branches with extreme differences using allele frequency (Shriver et al. 2004; Yi et al. 2010) or expression (Assis 2019; Jiang and Assis 2020) data. We applied these measurements here to identify particular analyses with strongly differentiated *P*-values (much higher or lower *P*-value) in one regression relative to the other two for a given gene. For example, a large 𝑑_AT_ value might reflect scenarios in which regression is strongly significant (*P*-value < 10^-6^) based on the amino acid tree, but not significant in the nucleotide or species tree-based regression (*P*-value > 0.05).

### Investigating a potential path forward with robust regression

Lastly, we explored the potential for a solution to the problem of tree mismatch with phylogenetic regression. In particular, we recently found promise in the application of robust estimators for improving the resistance of phylogenetic regression to evolutionary outliers (Adams et al. 2023). Given these findings, we sought to assess whether a robust estimator may yield better performance than the standard L2 estimator, which minimizes the mean squared error of predictions and is thus sensitive to outliers. To address this question, we employed the robust L1 estimator, which instead minimizes the mean absolute error of the predictions (Rousseeuw and Yohai 1984), helping to alleviate false positive rates associated with strong outliers by de-emphasizing large residuals compared to the L2 estimator. We applied L1-based regression to the same simulation conditions as before with 𝑛 ∈ {10,100,1000} species and varying levels of tree mismatch with birth rate parameter 𝜆 ∈ {10^-4^, 10^-3^, …, 10^2^} for known (simulated) trees, our case study that included gene tree estimation in addition to tree mismatch with 𝑛 = 10 species, and our empirical case study. We then compared false positive rates of L1-based regression to those obtained with L2-based regression.

## Results

### Illustrating the phylogenetic conundrum

We chose two simulation replicates to illustrate the phylogenetic conundrum for testing trait associations in the presence of tree discordance (Fig. 1). For these examples, we simulated a species tree 𝑆 and an associated gene tree 𝐺 given the multispecies coalescent process on *S*. We then simulated two statistically independent (𝛽 = 0) traits **x** and **y** using the species tree *S* (Fig. 1; top row), and separately using the gene tree *G* (Fig. 1; bottom row). Here we show two examples of matched phylogenetic regression in which the same tree was used to generate the trait data and analyze the trait data using PICs (Figs. 1c and d). Likewise, we provide two examples of mismatched phylogenetic regression in which the traits were generated according to the species tree, but the gene tree was incorrectly assumed when computing PICs (Fig. 1e), and the alternative scenario in which the traits were generated according to the gene tree, but the species tree was incorrectly assumed (Fig. 1f). *P*-values from matched phylogenetic regression SS (Fig. 1c) and GG (Fig. 1d) were not statistically significant, consistent with the independence of the two simulated traits (𝛽 = 0). However, both examples of mismatched phylogenetic regression based on SG (Fig. 1e) and GS (Fig. 1f) were statistically significant, providing evidence of a false positive association due to using the wrong tree for computing PICs. In these examples, the degree of false significance was higher for GS than for SG, as demonstrated by their *P*-values. Comparing the trait mappings provides some intuition, with evidence of hemiplasy when the response trait **y** is mapped to the incorrect tree (Figs. 1a and b).

### Impacts of tree mismatch on false positive rates of phylogenetic regression with simple architectures

Across our simulations, we found evidence of strong biases with incorrectly mismatched phylogenetic regression (GS and SG) compared to correctly matched regression (GG and SS; Figs. 2, S1, and S2). Specifically, false positive rates for GS and SG (red and orange) were higher than those for matched GG and SS (black), which yielded acceptable false positive rates of approximately 5% across all values of 𝜆 in our simulations (Figs. 2a-c). Thus, assuming the incorrect tree tended to mislead phylogenetic regression to reject the null hypothesis when the two traits were statistically independent (𝛽 = 0). The impact of phylogenetic mismatch was exacerbated with more species (Figs. 2a-c; top to bottom), shorter tree depths (Figs. 2, S1, and S2), and as the expected amount of ILS increased: false positive rate increased with speciation rate for both GS and SG (Figs. 2a-c; left to right in each panel). Specifically comparing the two mismatched scenarios (GS versus SG) revealed evidence of higher false positive rates for GS (red) than for SG (orange) across our simulations (Figs. 2, S1 and S2). That is, performance was worse when incorrectly assuming the species tree for traits generated from a gene tree (GS) than the reverse situation in which an incorrect gene tree was assumed for traits generated from a species tree (SG). The severity of false positive rate inflation was influenced by the overall depth of the species tree, with shorter tree depths exacerbating false positive rates comparatively (Fig. S1 vs. Fig. 2 vs. Fig. S2). Our results were similar when comparing false positive rates of mismatched regression with species trees generated via pure-birth and birth-death models (comparing Fig. 2 and Fig. S3). These findings were consistent with the overall distributions of *P*-values (Figs. 2d-i), and these impacts reflected topological distances between gene trees and species trees measured with the Robinson-Foulds metric (Figs. 2j-l), as well as probabilistic distances measured with the Hellinger distance (Figs. 2m-o).

### Complex architectures and mismatched phylogenetic regression

Our simulation study with more complex architectures for traits encoded by two, five, 10, or 100 loci continued to mirror results of simple, single tree simulations (Fig. 3). We found unacceptably high false positive rates across all tree sizes (10, 100, or 1000 species; rows in Fig. 3) and architectures (two, five, 10, or 100 loci; columns in Fig. 3) that were amplified with higher amounts of ILS (left to right on *x*-axes in Fig. 3) for analyses using the mismatched species tree and mismatched sets of gene trees. As with single tree regression (Fig. 2), increasing the sample size (i.e., increasing the number of species) only made the situation worse. Moreover, both scenarios of mismatch (i.e., mismatched species tree or mismatched set of gene trees; red and pink lines in Fig. 3) generated high false positive rates in many scenarios when compared to the appropriate false positive rate for analyses that assumed the correct set of gene trees (black lines; Fig. 3). However, incorrectly assuming a species tree tended to generate higher false positive rates than assuming an incorrect set of gene trees (red versus pink; Fig. 3). Increasing the architecture complexity (i.e., the number of loci encoding a trait; left to right columns in Fig. 3) suggested slight improvements, though false positive rates remained far higher than the typical 𝛼 = 0.05 cutoff for most scenarios. With large trees (100 or 1000 tips), even the smallest birth rates still exhibited remarkably high false positive rates, particularly for the mismatched species tree analyses. For example, false positive rates were estimated at ∼50% for 1000-tip trees with a birth rate of 𝜆 = 10^$&^(Fig. 3c).

**FIGURE 3.**
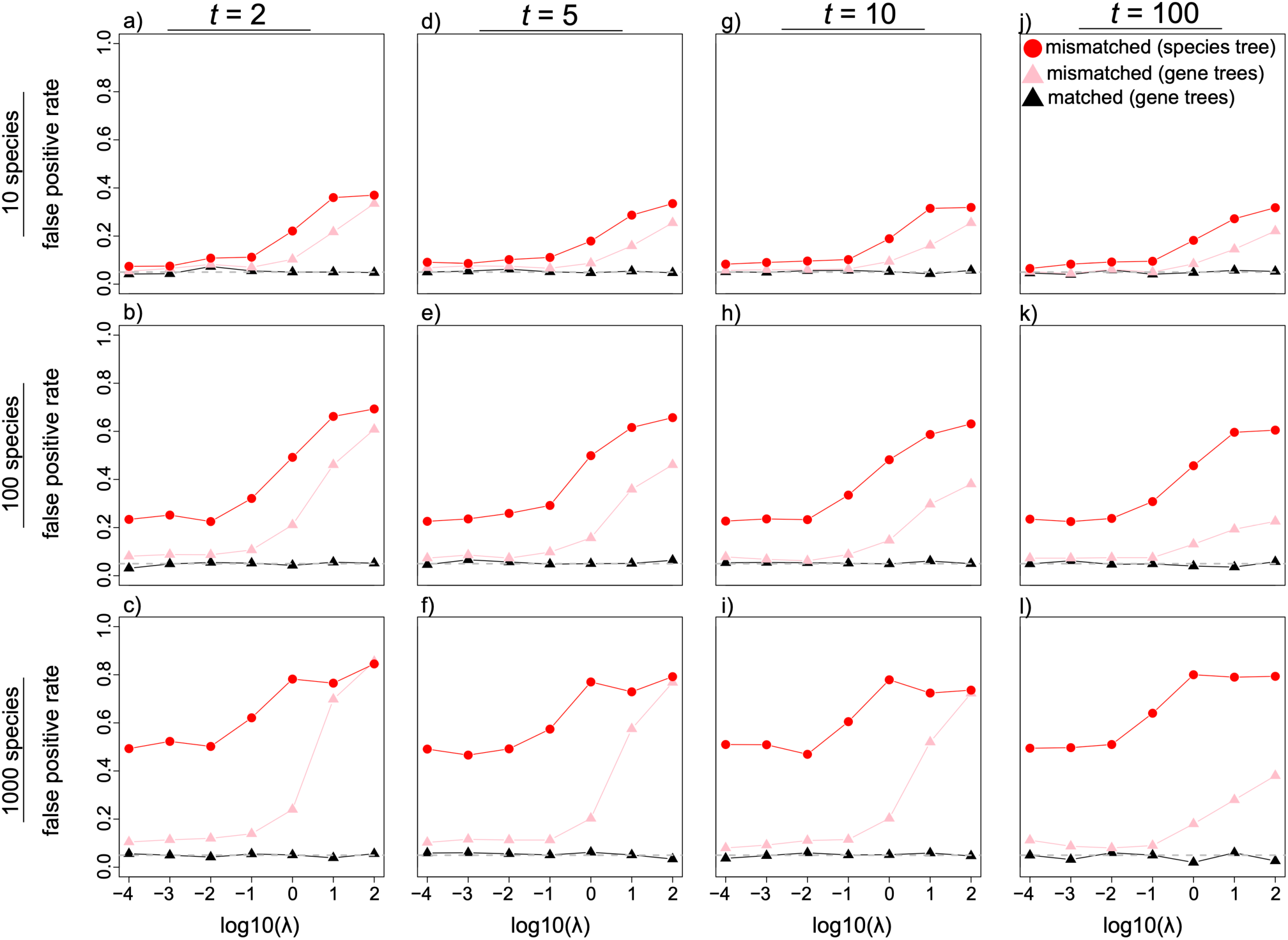
Tree mismatch misleads phylogenetic regression for traits with more complex architectures. Estimates of the false positive rates from simulations including 10 species (top row), 100 species (middle row), and 1000 species (bottom row) for birth-death simulations with birth rate 𝜆, death rate 𝜆/2, and root age of 10 coalescent units for mismatched species tree regression (red lines), mismatched gene tree regression (pink links), and matched gene tree sets (black lines). Results shown for traits encoded by two loci (a-c), five loci (d-f), 10 loci (g-i), and 100 loci (j-l). The two traits were statistically independent (𝛽 = 0) for all simulations. Horizontal dashed lines mark the commonly used false positive rate of 𝛼 = 0.05.

### Simulation case study: phylogenetic estimation error and tree mismatch together

Analyses conducted on known trees can be markedly different from those based on estimated trees. Regardless of whether known (Fig. 4a) or estimated (Fig. 4b) trees are assumed, mismatched regression can amplify false positive rates. Perhaps expectedly, we found higher false positive rates when using estimated versus known phylogenies in many cases. We also found that estimation error can increase false positive rates for matched GG and SS phylogenetic regression (black lines; Fig. 4). The effects of estimation error on these matched scenarios were still less pronounced than those on mismatched GS and SG (red and orange lines; Fig. 4), however. Increasing the speciation rate tended to exacerbate false positive rates for all GG, SS, GS, and SG scenarios with estimated trees (Fig. 4b), whereas known matched regression scenarios (GG and SS) were unaffected (Fig. 4a). Comparing differences between log-scaled *P*-values of known and estimated trees further highlighted these findings (Fig. 4c), with the largest differences in significance between known and estimated analyses observed in the matched SS, followed by SG, GG, and GS. This result likely reflects the higher relative false positive rates for SS when using estimated versus known trees (black lines; Fig. 4b), whereas known matched analyses demonstrate acceptable false positive rates of 0.05 (black lines; Fig. 4a). In these scenarios, phylogenetic regression with GS and SG were strongly influenced by tree mismatch with known trees and estimated trees (red and orange lines in Figs. 4a and b).

**FIGURE 4.**
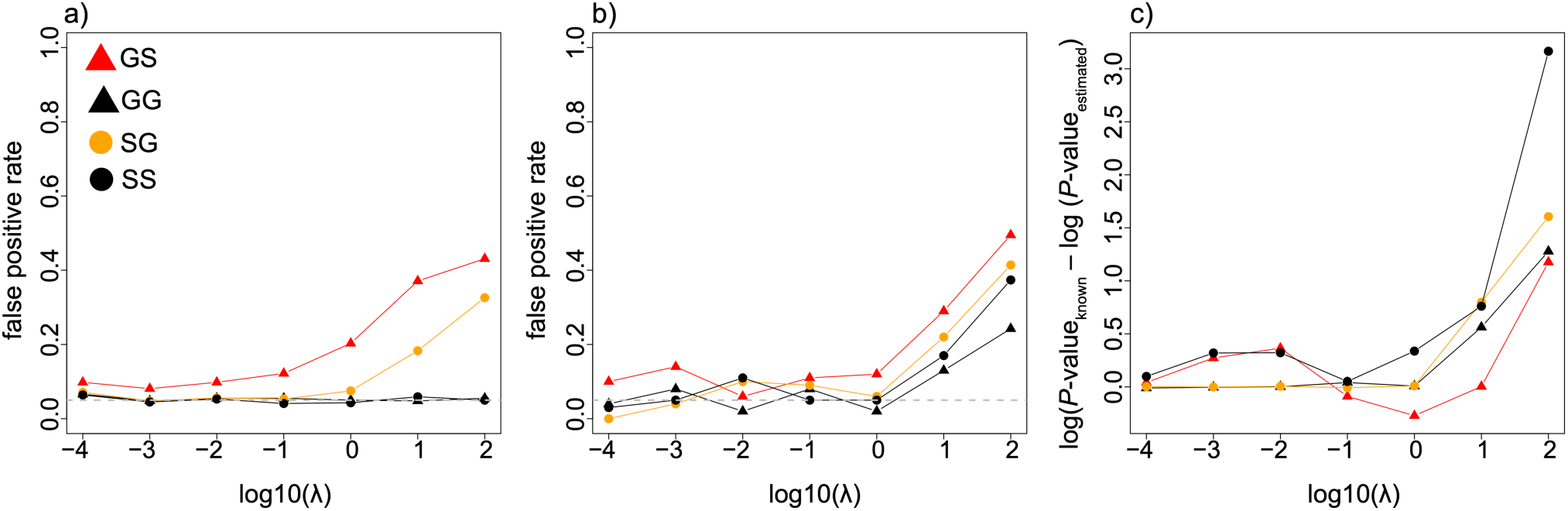
Case studying the impacts of both tree mismatch and tree estimation error on phylogenetic regression under birth-death simulations with birth rate 𝜆, death rate 𝜆/2, and root age of 10 coalescent units. Depicted are false positive rates of the two mismatched scenarios (GS and SG) and the two matched scenarios (GG and SS) when known trees were assumed for regression (a) and alternatively, when estimated trees were assumed for regression (b) for 𝑛 = 10 species. Difference in log-scaled *P*-value between the regression model fit to the known tree and to the estimated tree (c).

### Tree mismatch and statistical power of phylogenetic regression

Next, we evaluated the potential for phylogenetic mismatch to influence the statistical power of regression when two traits are indeed statistically associated (𝛽 > 0). When compared with false positive rates, the effects of mismatched trees on power appear to be less dramatic and fluctuate depending on the value of 𝛽 and number of species (Fig. 5). In many cases, however, we found evidence that mismatched regression can decrease statistical power. This finding is perhaps most apparent in our simulations with 𝛽 = 0.50 and 100 species (Fig. 5e), as well as with 𝛽 = 0.25 and 1000 species (Fig. 5c), in which mismatched GS scenarios demonstrated comparatively lower power (red lines; Fig. 5). Mismatched SG scenarios also exhibited lower power than matched regression in some examples (orange versus black; Fig. 5). However, impacts were less apparent for smaller trees (10 species; top row Fig. 5) and deeper species trees (Fig. S4 vs. Fig. 5 vs. Fig. S5).

**FIGURE 5.**
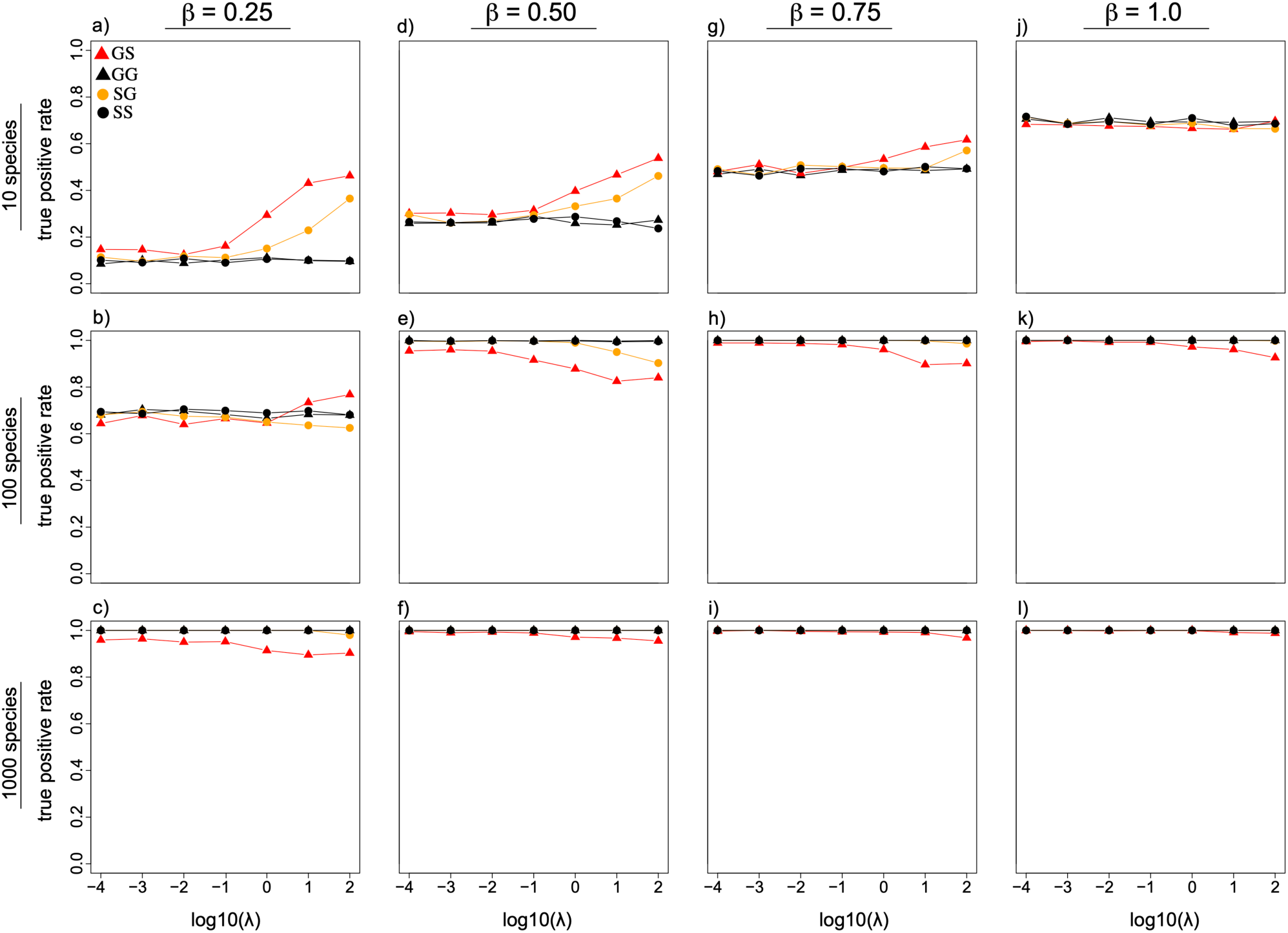
Impacts of mismatched phylogenetic regression on statistical power to detect true trait associations. Estimates of true positive rates for 10 species (top row), 100 species (middle row), and 1000 species (bottom row) for birth-death simulations with birth rate 𝜆, death rate 𝜆/2, and root age of 10 coalescent units. Results are shown for 𝛽 = 0.25 (a-c), 𝛽 = 0.50 (d-f), 𝛽 = 0.75 (g-i), and 𝛽 = 1.0 (j-l).

### Empirical case study: investigating phylogenetic mismatch and gene expression data

Most apparent in our exploration of phylogenetic regression using mammalian gene expression data are the differences in significance levels among analyses using the species tree, nucleotide gene tree, and amino acid gene tree (Figs. 6a-c). For instance, in heart tissue, we observed the smallest number of outliers (red points) for the 𝑑_ST_ divergence statistic corresponding to analyses using the species tree (Fig. 6a; 38 genes), followed by the 𝑑_NT_ statistic for analyses using the nucleotide gene tree (Fig. 6b; 107 genes), and finally by the 𝑑_AT_ statistic for analyses using the amino acid gene tree (Fig. 6c; 643 genes). Thus, using the amino acid gene tree for phylogenetic regression resulted in the identification of many more significantly associated genes in female and male heart tissue than using either the species tree or nucleotide gene tree.

**FIGURE 6.**
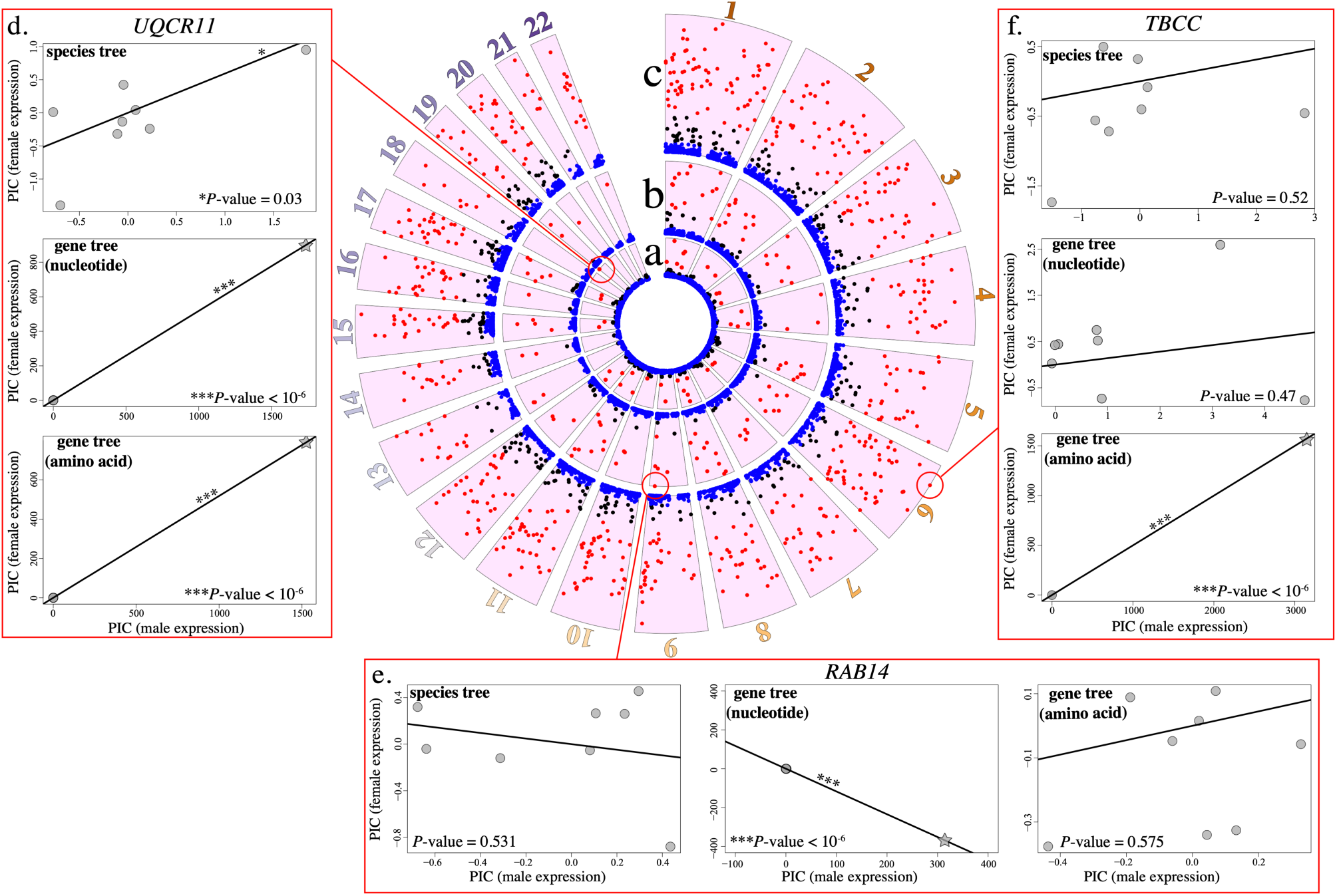
Exploring phylogenetic regression and mismatch for a mammalian gene expression dataset with divergence statistics. Results shown across 22 autosomes for heart tissue expression measurements of 4,068 genes with measurable expression across species, with computed distance statistics 𝑑_ST_(inner track a), 𝑑_NT_(middle track b), and 𝑑_AT_(outer track c) based on L2-based phylogenetic regression. Empirical case studies comparing phylogenetic regression based on the species tree (d), nucleotide gene tree (e), and amino acid gene tree (f) are shown for analyses with anomalously high 𝑑_ST_, 𝑑_NT_, and 𝑑_AT_, respectively. Colors of points in circos plot (a-c) indicate relative level of divergent *P*-values, with blue indicating not significant, black indicating *P*-value < 0.05, and red indicating strong outliers with *P*-value < 1.229 × 10^$4^ after applying Bonferroni correction (Bonferroni 1936). Points depicted as gray stars indicate evidence of singular phylogenetic outliers found in specific analyses (d-f).

These genome-level explorations also allowed us to identify the largest outlier genes based on their values of 𝑑_ST_, 𝑑_NT_, and 𝑑_AT_ (Fig. 6d-f). The largest outlier based on 𝑑_ST_ was *UQCR11* (Fig. 6d), which is involved in the mitochondrial electron transport chain. All three analyses revealed a positive relationship between female and male expression in this gene, though with much weaker significance when using the species tree than when using either of the two gene trees (Fig. 6d). The largest outlier based on 𝑑_NT_ was *RAB14* (Fig. 6e), which is involved in intracellular membrane trafficking. For this gene, the nucleotide tree yielded a highly significant negative relationship between female and male expression, whereas the other two trees did not produce significant results (Fig. 6e). Finally, the largest outlier based on 𝑑_AT_ was *TBCC* (Fig. 6f), which is one of four genes involved in the pathway leading to correctly folded beta-tubulin from folding intermediates. For this gene, the amino acid tree regression produced a highly significant positive relationship between female and male expression, whereas the other two analyses did not yield a significant association (Fig. 6f).

Next, we evaluated overlap in statistically significant genes estimated using the three regression strategies (i.e., assuming the species tree, nucleotide gene tree, or amino acid gene tree) across tissues. We first considered the fraction of statistically significant (*P*-value < 0.05) analyses from a tissue-level perspective. For each of three tissues considered (brain, heart, and kidney), we found substantial overlap in the percentages of genes with estimates of statistically significant relationships between female and male expression (Fig. 7). That is, within a given tissue, the total fractions of significant genes were similar for regression based on the species tree, amino acid gene tree, and nucleotide gene tree. However, consistent with our previous findings in heart (Figs. 7a-c), phylogenetic regression based on the amino acid gene tree yielded the largest percentage of uniquely significant genes for all tissues, with 24, 23, and 22% significant for brain, heart, and kidney analyses, respectively (Fig. 7). Given these results, we then computed log-likelihoods of the fitted phylogenetic regression model for the three tissues and the three strategies. All three tissues agree that model fit was highest on average when assuming the species tree, followed by the nucleotide gene tree, and finally the amino acid gene tree (Fig. 8). That is, phylogenetic regression models tend to fit the species tree best and the amino acid gene tree worst, with the nucleotide gene tree fit representing an intermediate between the two. Thus, perhaps the excess of uniquely significant genes identified by regression using the amino acid gene tree is due to a poor fit.

**FIGURE 7.**
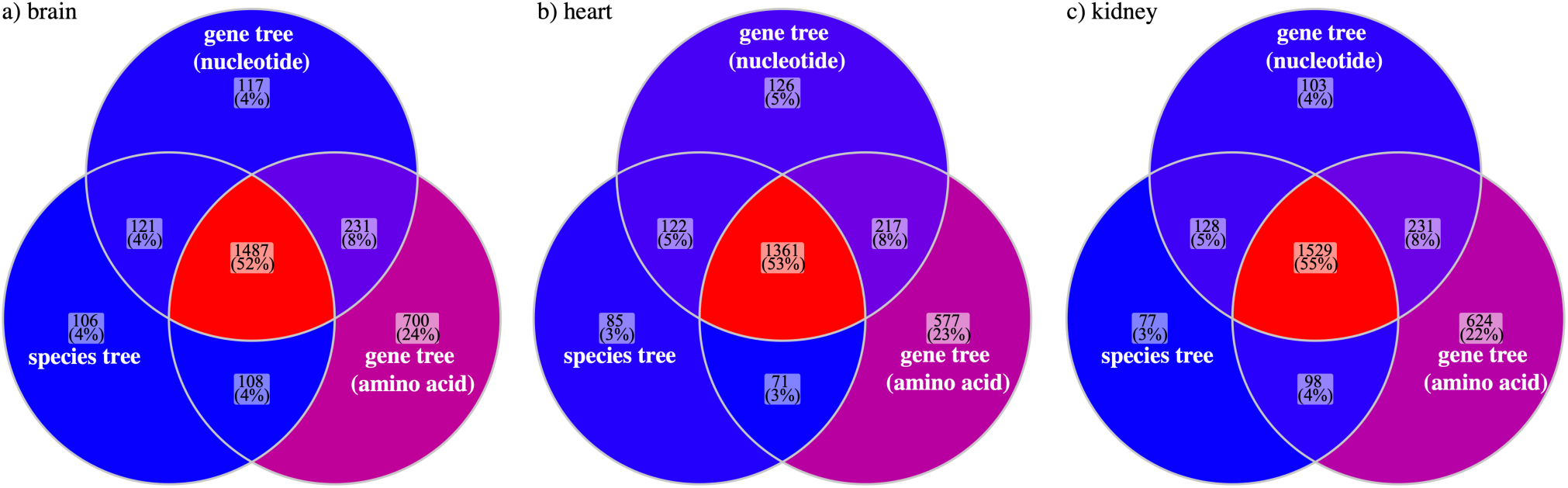
Venn diagrams displaying the percentage of overlap in statistically significant genes for brain (a), heart (b), and kidney (c) expression levels in a mammalian dataset based on phylogenetic regression applied by assuming the species tree (left circles), nucleotide gene tree (top circles), or amino acid gene tree (right circles). Colors indicate the relative percentage of statistically significant genes across analyses.

**FIGURE 8.**
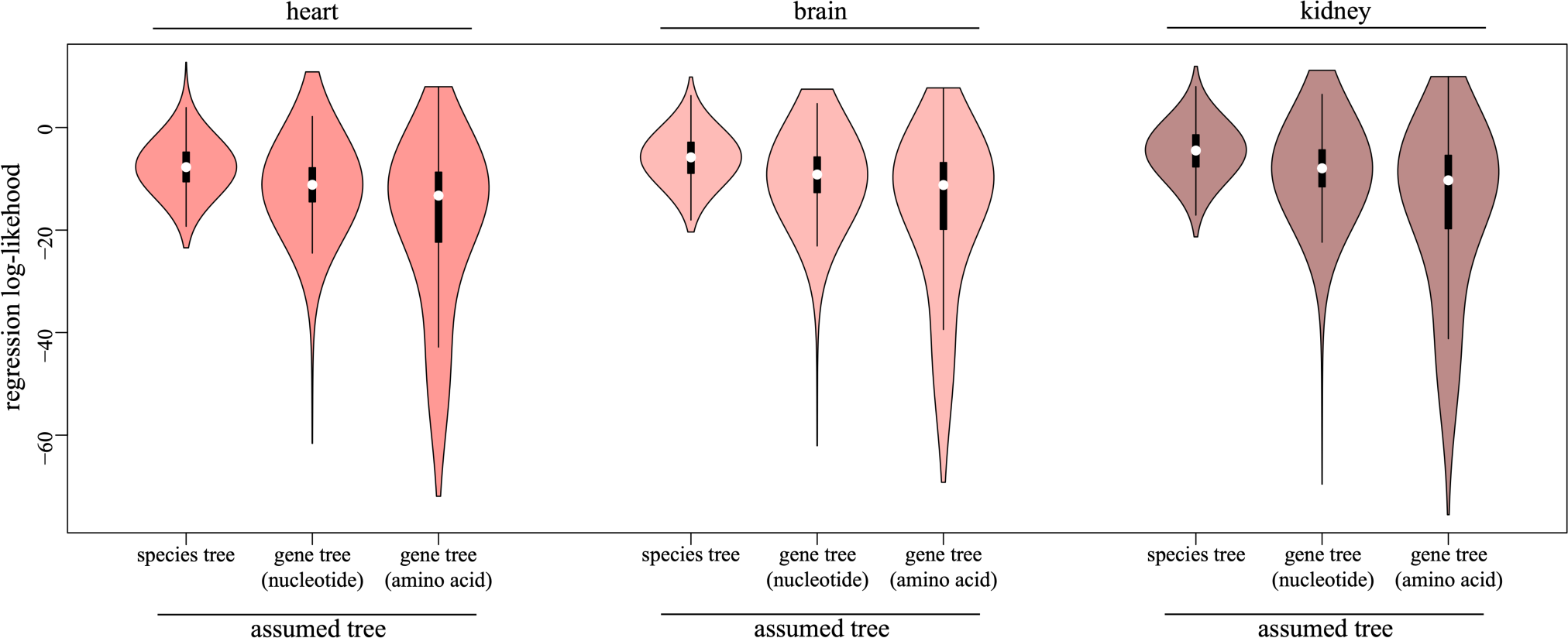
Violin plots summarizing the distributions of model fit measured by log-likelihood for phylogenetic regression applied to gene expression from a mammalian dataset. Results shown across tissues (heart, brain, and kidney) and the three regression strategies that assume either the species tree, nucleotide gene tree, or amino acid gene tree.

### Exploring the potential for robust phylogenetic regression

Lastly, we explored the application of robust regression in the context of tree mismatch. We found evidence that robust L1-based regression can reduce false positive rates, at least compared to analyses that use standard L2-based regression for both known (Fig. 9a and b) and estimated phylogenies (Fig. 9c). In particular, L1-based regression yielded comparatively fewer false positives for GS (solid versus dashed red lines; Figs. 9) and SG (solid versus dashed orange lines; Figs. 9) under most conditions of mismatched regression. When considering our analyses of simulations that used estimated rather than known trees with 𝑛 = 10 species, we still found relatively lower false positive rates when using L1-versus L2-based regression (Fig. 9c), albeit to a lesser degree. Reflecting on our empirical case studies, we found several interesting differences between robust L1-based and standard L2-based regression when assuming the species tree, nucleotide gene tree, or amino acid gene tree (Fig. 10). In several examples, L1-based regression yielded comparatively smaller *P*-values (i.e., higher significance), sometimes leading to the inferences of statistically significant relationships not identified by L2-based regression (Figs. 10a-c). In others, L1-based regression returned comparatively larger *P*-values, such that it did not infer statistically significant relationships found with L2-based regression (Figs. 10d-f).

**FIGURE 9.**
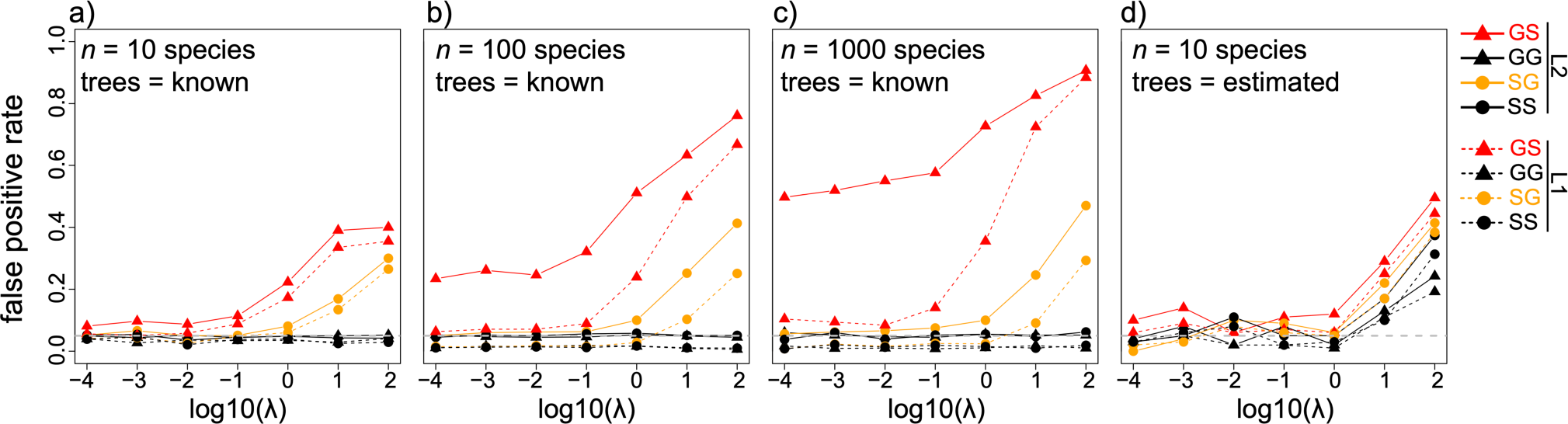
Can robust estimators help with tree mismatch? Results showing estimated false positive rates when using known trees with 10 species (a), 100 species (b), and 1000 species (c) for robust L1-based regression (dashed lines) alongside standard L2-based regression (solid lines) under birth-death simulations with birth rate 𝜆, death rate 𝜆/2, and root age of 10 coalescent units. Estimated false positive rates are also shown for L1- and L2-based regression with estimated trees for our simulation case study with 𝑛 = 10 species (d). Horizontal solid gray lines mark the typically accepted false positive rate of 0.05.

**FIGURE 10.**
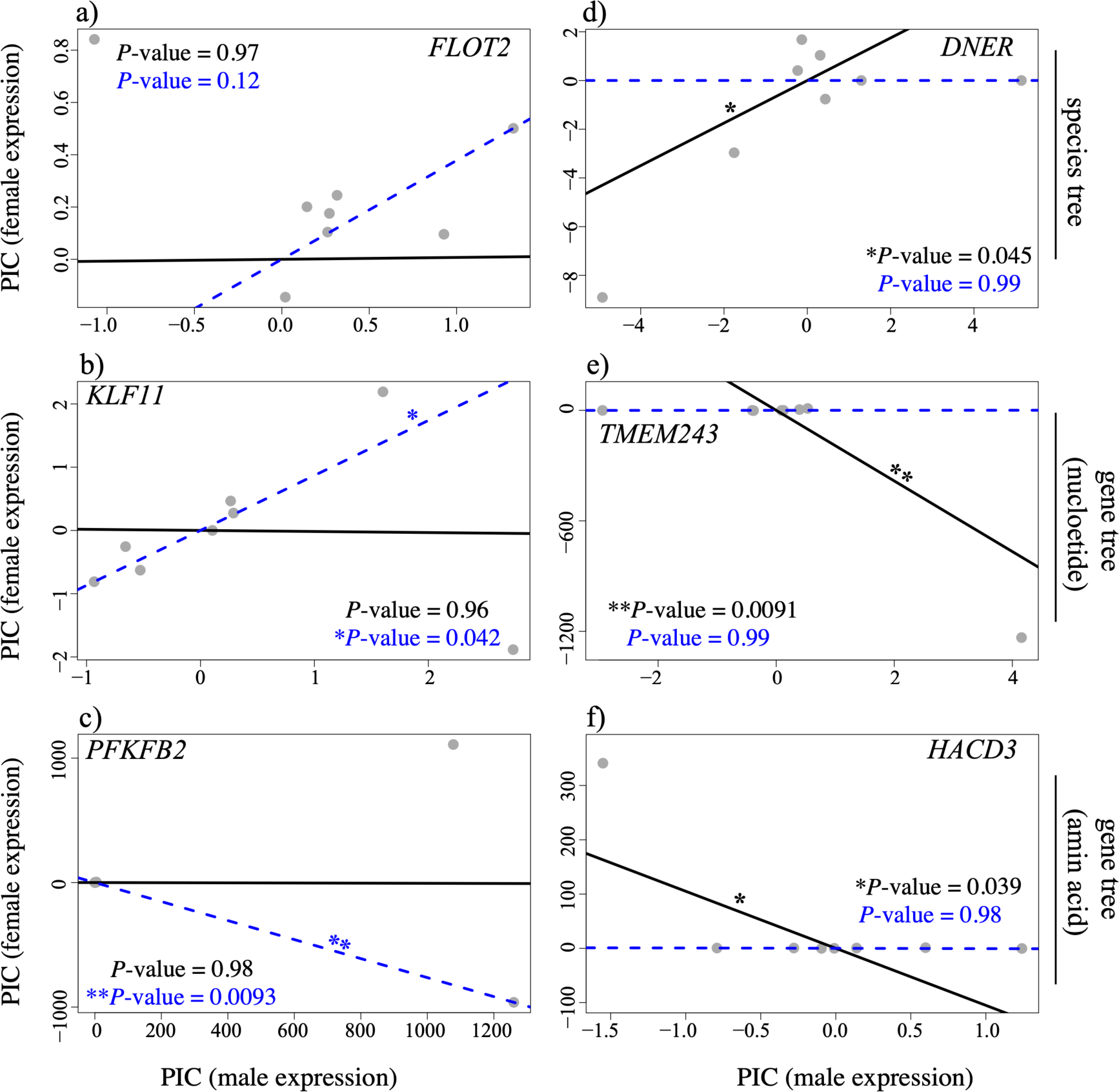
Can robust estimators help with tree mismatch? Empirical examples from the mammalian gene expression data showing interesting differences in *P*-values obtained with standard L2-based (black lines) and robust L1-based (blue dashed lines) linear regression using the species tree (top row), nucleotide gene trees (middle row), and amino acid gene trees (bottom row).

## Discussion

A choice of trees is always required when conducting phylogenetic regression. Yet deciding on a particular tree or trees is often difficult and unlikely to get easier anytime soon. Collectively, our analyses underscore these challenges and expand our understanding of statistical pitfalls when trees are mismatched for testing relationships between traits with phylogenetic regression. To summarize, assuming the wrong tree may lead us to overestimate associations between traits that are truly independent—regardless of whether considering shallow or deep trees, few or many species, simple or complex architectures, and known or estimated trees. That is, the choice of tree(s) matters. Our study is the first to present these findings for phylogenetic regression, and thus we focused on ILS-based mismatch resulting from gene tree-species tree discordance—a topic that has held our field captive for decades. Examining other potential sources of conflict (e.g., recombination, selection, and introgression) is arguably a worthwhile next step as it will incorporate other realistic processes that are often encountered in empirical data. Moreover, future studies with expanded simulations will help us better understand the simultaneous effects of tree mismatch and estimation error (i.e., expanding results shown in Fig. 4), though it is worth emphasizing the computationally intensive and expensive demands of these multi-layered analyses that span simulation and inference of species trees, gene trees, sequence alignments, and trait evolution.

Mirroring similar types of evolutionary analyses (Hahn and Nakhleh 2016b; Mendes and Hahn 2016; Guerrero and Hahn 2018; Mendes et al. 2018, 2019; Hibbins et al. 2020; Hibbins et al. 2022), tests of trait associations are sensitive to gene tree-species tree conflict. Phylogenetic mismatch due to ILS amplified false positive rates compared to properly matched analyses in many conditions. The rate of speciation was an important factor in determining the degree of severity, with faster rates (𝜆 > 10^$0^) tending to yield higher false positive rates. Because the expected length of internal branches (i.e., time between speciation events) is inversely related with speciation rate, faster speciation rates yield shorter internal branches, which in turn can amplify phylogenetic conflict and ILS by providing less time for coalescent events in ancestral branches. In mismatched analyses (GS and SG), the degree of severity also reflected the distance between the true and assumed tree, such that larger Robinson-Foulds and Hellinger distances were associated with higher ILS and higher false positive rates. Increasing the sample size (i.e., increasing the number of species) only exacerbated these issues, and yet small trees were certainly not immune. Thus, scenarios with rampant ILS will likely prove the most challenging when applying phylogenetic regression. When compared with estimates of false positive rates, statistical power to detect real trait associations appeared less affected by tree mismatch, though future studies will be needed to better understand some of the patterns we uncovered.

Another surprising trend emerged when specifically comparing the two scenarios of tree mismatch—higher false positive rates with GS than with SG. That is, incorrectly assuming a species tree for traits simulated under a gene tree tended to be worse than the opposite scenario. Neither mismatched scenario performed well for high speciation rates, and yet our findings suggest that assuming an incorrect gene tree may represent a potential lesser of two evils, at least in these scenarios explored here. Dissecting this pattern further by comparing mean magnitudes of PICs for tree cherries (i.e., nodes with exactly two extant descendants) provided evidence of increasingly larger contrasts for GS than GG regression with speciation rate (Fig. S6). This pattern was consistent in our simulation case study that also included phylogenetic estimation error. In an ideal world, one would always match the tree to the trait perfectly (i.e., GG and SS), but this is neither always possible nor probable. Moreover, our simulations with complex architectures continued these results: incorrectly assuming the species tree nearly always amplified false positive rates, as did assuming an incorrect set of gene trees that were unrelated to the traits of interest.

Clearly, the reliability of an assumed tree is likely to be a major determinant of the reliability of an evolutionary hypothesis test. When designing this study, we first focused on using known trees to isolate and understand the behavior of mismatched regression due to ILS alone. We then realized that we needed to consider an important elephant in the room: in practice, phylogenetic regression is conducted using estimated trees rather than known trees. Of course, we seldom (if ever) estimate a phylogeny to perfection, and our findings argue for increased vigilance against both tree mismatch and estimation error, as even matched regression suffered in the presence of gene tree estimation error. Our study represents the first to investigate the impacts of ILS-based tree mismatch on phylogenetic regression, and thus, we used scenarios of both known and estimated trees to dissect their impacts separately, as well as considered them together for smaller trees. Using estimated trees tended to yield poorer performance, but this observation is typically the reality for empirical studies that must rely on estimated trees. Future simulation studies seeking to fully explore the scope and scale of both tree mismatch and estimation error are likely to be quite computationally costly, requiring multiple levels of analyses that begin with species tree simulation, to gene tree simulation, sequence alignment simulation, and character trait simulation, and then followed by gene tree estimation, species tree estimation, and finally, phylogenetic regression using different trees of various sizes and shapes. Though focused for computational feasibility, our simulations nonetheless argue that both mismatch and estimation error are important, and we found evidence of alarming biases in the presence of both. Building on our simulation-based investigations, we sought to use an expansive gene expression dataset to study the potential impacts of regression strategies that assume different trees in empirical analyses. Because gene sequence and expression divergence are correlated (Duret and Mouchiroud 2000; Pál et al. 2001; Subramanian and Kumar 2004; Lemos et al. 2005; Assis and Kondrashov 2014), gene expression is typically assumed to evolve more or less according to an associated local gene tree. Thus, we used these empirical data as a best-case scenario in which one might have a fighting chance to match tree with trait. Yet clearly tree choice matters even in these scenarios. Our simulations of both simple and complex architectures complement these findings, which are broadly relevant for many studies that explicitly assume only a single tree for admittedly complex traits (e.g., Ross et al. 2004; Al-Kahtani et al. 2004; Kamilar and Cooper 2013; Gu 2016; Dunn et al. 2018; Chen et al. 2023). Perhaps most apparent in these analyses is the potential for stark differences in regression significance depending on the assumed tree. Further, our study suggests that using amino acid gene trees often yields strikingly different results than assuming either the species tree or nucleotide gene trees. In light of our simulation study with estimated trees, these results together indicate that gene tree estimation error can be a major driver of false significance in trait association studies. Altogether, our findings underscore the difficulties of phylogenetic regression in the presence of tree uncertainty and call for increased vigilance against phylogenetic mismatch—whether due to ILS, estimation error, or other sources.

It is often much easier to find flaws than to find solutions; seldom is it satisfying to simply point out major issues without offering at least a hope of a remedy. We found that to be the case here. While the primary purpose of this study was to provide a first perspective on the dangers of tree mismatch for regression, we also explored a potential path forward with the application of robust estimators for phylogenetic regression. Use of robust phylogenetic regression improved inferences for both known and estimated trees. Additionally, we illustrated several examples of large differences between *P*-values obtained with L1- and L2-based regression, most of which altered conclusions about tested relationships. Altogether, our analyses suggest that robust estimators might provide a potential, albeit imperfect, solution to issues raised by tree mismatch. We can say with confidence that robust phylogenetic regression was never meant to be a panacea for all ailments that might afflict PCMs. Progress—not perfection—is the goal, and more studies are needed to explore the possibilities and space of phylogenetic mismatch and the potential for different types of robust estimators with different types of model violations. Strategies that fit Pagel’s lambda (Pagel 1999b) to raw regression residuals to estimate the phylogenetic variance-covariance matrix may also prove helpful for modeling traits on discordant gene trees (i.e., mixing findings of Mazel et al 2016 and Mendes et al. 2019). Future studies that employ both robust estimators and other recent advances in phylogenetic modeling (e.g., phylogenomic comparative methods; Hibbins et al. 2023) may prove helpful in this context.

## Funding And Acknowledgements

This work was supported by National Science Foundation grants DBI-2130666, DEB-2302258, DEB-2392257, and BCS-2001063, National Institutes of Health grants R35GM142438 and R35GM128590, and start-up funds provided by the University of Arkansas. This research was also supported by the Arkansas High Performance Computing Center, which is funded through multiple National Science Foundation grants and the Arkansas Economic Development Commission, as well as Research Computing at Florida Atlantic University.

## Disclosure Statement

The authors of this study do not report any conflicts of interest.

## Notes

### Competing Interest Statement

The authors have declared no competing interest.

## References

Adams D.C. 2013. Comparing evolutionary rates for different phenotypic traits on a phylogeny using likelihood. Syst Biol. 62:181–192.

Adams R.H., Blackmon H., DeGiorgio M., 2021. Of traits and trees: probabilistic distances under continuous trait models for dissecting the interplay among phylogeny, model, and data. Syst Biol. 70:660–680.

Adams R.H., Blackmon H., Reyes-Velasco J., Schield D.R., Card D.C., Andrew A.L., Waynewood, N, Castoe, T.A. 2016. Microsatellite landscape evolutionary dynamics across 450 million years of vertebrate genome evolution. Genome 59:295–310.

Adams R.H., Cain Z., Assis R., DeGiorgio, M. 2023. Robust phylogenetic regression. bioRxiv. 2022– 08.

Adams R.H., Schield D.R., Card D.C., Castoe T.A. 2018. Assessing the impacts of positive selection on coalescent-based species tree estimation and species delimitation. Syst Biol. 67:1076–1090.

Al-Kahtani M.A., Zuleta C., Caviedes-Vidal E., Garland, Jr, T., 2004. Kidney mass and relative medullary thickness of rodents in relation to habitat, body size, and phylogeny. Physiological and Biochemical Zoology. 77:346–365.

Assis R. 2019. Lineage-specific expression divergence in grasses is associated with male reproduction, host-pathogen defense, and domestication. Genome Biol Evol. 11:207–219.

Assis R., Kondrashov A.S. 2014. Conserved proteins are fragile. Mol. Biol. Evol. 31:419–424.

Avise J.C., Robinson T.J. 2008. Hemiplasy: a new term in the lexicon of phylogenetics. Syst Biol. 57:503–507.

Bastide P., Ho L.S.T., Baele G., Lemey P., Suchard, M.A.. 2021. Efficient Bayesian inference of general Gaussian models on large phylogenetic trees. The Annals of Applied Statistics. 15: 971–997.

Bastide P., Soneson C., Stern D.B., Lespinet O., Gallopin M. 2023. A Phylogenetic Framework to Simulate Synthetic Interspecies RNA-Seq Data. Mol Biol Evol. 40: 269.

Beaulieu J.M., Jhwueng D., Boettiger C., O’Meara B.C. 2012. Modeling stabilizing selection: expanding the Ornstein–Uhlenbeck model of adaptive evolution. Evolution. 66:2369–2383.

Bertram J., Fulton B., Tourigny J., Pena-Garcia Y., Moyle L.C., Hahn M.W. 2022. CAGEE: computational analysis of gene expression evolution. bioRxiv.:2011–2022.

Blomberg S.P., Garland Jr T., Ives A.R. 2003a. Testing for phylogenetic signal in comparative data: behavioral traits are more labile. Evolution. 57:717–745.

Blomberg S.P., Lefevre J.G., Wells J.A., Waterhouse M. 2012. Independent contrasts and PGLS regression estimators are equivalent. Syst Biol. 61:382–391.

Bonferroni C. 1936. Teoria statistica delle classi e calcolo delle probabilita. Pubbl. del R Ist. Super. di Sci. Econ. e Commericiali di Firenze. 8:3–62.

Borges R., Boussau B., Szöllősi G.J., Kosiol C. 2020. Pervasive selection biases inferences of the species tree. bioRxiv.:2007–2020.

Brawand D., Wagner C.E., Li Y.I., Malinsky M., Keller I., Fan S., Simakov O., Ng A.Y., Lim Z.W., Bezault E. 2014. The genomic substrate for adaptive radiation in African cichlid fish. Nature. 513:375–381.

Cavalli-Sforza L.L., Edwards A.W.F. 1967. Phylogenetic analysis. Models and estimation procedures. Am J Hum Genet. 19:233.

Chen F., Li Z., Zhang X., Wu P., Yang W., Yang J., Chen X., Yang J.R., 2023. Phylogenetic comparative analysis of single-cell transcriptomes reveals constrained accumulation of gene expression heterogeneity during clonal expansion. Mol Biol and Evol. 40:5

Chen J., Swofford R., Johnson J., Cummings B.B., Rogel N., Lindblad-Toh K., Haerty W., Di Palma F., Regev A. 2019. A quantitative framework for characterizing the evolutionary history of mammalian gene expression. Genome Res. 29:53–63.

Copetti D., Búrquez A., Bustamante E., Charboneau J.L.M., Childs K.L., Eguiarte L.E., Lee S., Liu T.L., McMahon M.M., Whiteman N.K. 2017. Extensive gene tree discordance and hemiplasy shaped the genomes of North American columnar cacti. Proceedings of the National Academy of Sciences. 114:12003–12008.

DeGiorgio, M., Rosenberg, N.A., 2016. Consistency and inconsistency of consensus methods for inferring species trees from gene trees in the presence of ancestral population structure. Theor. Pop. Bio. 110:12–24.

Degnan J.H., Rosenberg N.A. 2009. Gene tree discordance, phylogenetic inference and the multispecies coalescent. Trends Ecol Evol. 24:332–340.

Diaz-Uriarte R., Garland Jr T. 1996. Testing hypotheses of correlated evolution using phylogenetically independent contrasts: sensitivity to deviations from Brownian motion. Syst Biol. 45:27–47.

Diaz-Uriarte R., Garland Jr T. 1998. Effects of branch length errors on the performance of phylogenetically independent contrasts. Syst Biol. 47:654–672.

Dimayacyac J.R., Wu S., Pennell M. 2023. Evaluating the Performance of Widely Used Phylogenetic Models for Gene Expression Evolution. bioRxiv.:2002–2023.

Doña J., Johnson, K.P. 2023. Host body size, not host population size, predicts genome-wide effective population size of parasites. Evolution Letters.7:285–292.

Dunn C.W., Zapata F., Munro C., Siebert S., Hejnol, A., 2018. Pairwise comparisons across species are problematic when analyzing functional genomic data. Proceedings of the National Academy of Sciences. 115:E409–E417.

Duret L, Mouchiroud D. 2000. Determinants of substitution rates in mammalian genes: expression pattern affects selection intensity but not mutation rate. Mol. Biol. Evol.17: 68–070.

Edwards S. V. 2009. Is a new and general theory of molecular systematics emerging? Evolution (N Y). 63:1–19.

Einum S., Fleming I.A. 2007. Of chickens and eggs: diverging propagule size of iteroparous and semelparous organisms. Evolution. 61:232–238.

Felenstein J. 2004. Inferring phylogenies. Sinauer associates Sunderland, MA.

Felsenstein J. 1973. Maximum-likelihood estimation of evolutionary trees from continuous characters. Am J Hum Genet. 25:471.

Felsenstein J. 1985. Phylogenies and the comparative method. Am Nat. 125:1–15.

Francis W.R., Canfield D.E. 2020. Very few sites can reshape the inferred phylogenetic tree. PeerJ. 8:e8865.

Gardner J.D., Organ C.L. 2021. Evolutionary Sample Size and Consilience in Phylogenetic Comparative Analysis. Syst Biol.

Garland Theodore J., Ives A.R. 2000. Using the past to predict the present: confidence intervals for regression equations in phylogenetic comparative methods. Am Nat. 155:346–364.

Grafen A. 1989. The phylogenetic regression. Philosophical Transactions of the Royal Society of London. B, Biological Sciences. 326:119–157.

Gu X. 2016. Understanding tissue expression evolution: from expression phylogeny to phylogenetic network. Briefings in bioinf. 17:249–254.

Guerrero R.F., Hahn M.W. 2018. Quantifying the risk of hemiplasy in phylogenetic inference. Proceedings of the National Academy of Sciences. 115:12787–12792.

Hahn M.W., Nakhleh L. 2016a. Irrational exuberance for resolved species trees. Evolution. 70:7–17.

Hansen T.F. 1997. Stabilizing selection and the comparative analysis of adaptation. Evolution (N Y). 51:1341–1351.

Harmon L.J., Losos J.B., Jonathan Davies T., Gillespie R.G., Gittleman J.L., Bryan Jennings W., Kozak K.H., McPeek M.A., Moreno-Roark F., Near T.J. 2010. Early bursts of body size and shape evolution are rare in comparative data. Evolution. 64:2385–2396.

Harvey P.H., Pagel M.D. 1991a. The comparative method in evolutionary biology. Oxford university press Oxford.

Hibbins M.S., Breithaupt L.C., Hahn M.W., 2023. Phylogenomic comparative methods: Accurate evolutionary inferences in the presence of gene tree discordance. Proceedings of the National Academy of Sciences.120:22.

He C., Liang D., Zhang P. 2020. Asymmetric distribution of gene trees can arise under purifying selection if differences in population size exist. Mol Biol Evol. 37:881–892.

Hensen N., Bonometti L., Westerberg I., Brännström I.O., Guillou S., Cros-Aarteil S., Calhoun S., Haridas S., Kuo A., Mondo S., Pangilinan J. 2023. Genome-scale phylogeny and comparative genomics of the fungal order Sordariales. Molecular Phylogenetics and Evolution. 189:107938.

Hibbins M.S., Breithaupt L.C., Hahn M.W. 2023. Phylogenomic comparative methods: Accurate evolutionary inferences in the presence of gene tree discordance. Proceedings of the National Academy of Sciences. 120:e2220389120.

Hibbins M.S., Gibson M.J.S., Hahn M.W. 2020. Determining the probability of hemiplasy in the presence of incomplete lineage sorting and introgression. Elife. 9:e63753.

Hobolth A., Dutheil J.Y., Hawks J., Schierup M.H., Mailund T. 2011a. Incomplete lineage sorting patterns among human, chimpanzee, and orangutan suggest recent orangutan speciation and widespread selection. Genome Res. 21:349–356.

Huey R.B., Garland Jr T., Turelli M. 2019. Revisiting a key innovation in evolutionary biology: Felsenstein’s “phylogenies and the comparative method.” Am Nat. 193:755–772.

Jeschke J.M., Kokko H. 2009. The roles of body size and phylogeny in fast and slow life histories. Evolutionary Ecology. 23:867–878.

Jiang X, Assis R. 2020. Population-specific genetic and expression differentiation in Europeans. Genome Biol Evol. 12:358–69.

Kamilar J.M., Cooper N. 2013. Phylogenetic signal in primate behaviour, ecology and life history. Philosophical Transactions of the Royal Society B: Biological Sciences. 368:20120341.

Koch H., DeGiorgio M. 2020. Maximum Likelihood estimation of species trees from gene trees in the presence of ancestral population structure. Genome Biol Evol. 12:3977–3995.

Kubatko L.S., Degnan J.H. 2007. Inconsistency of phylogenetic estimates from concatenated data under coalescence. Syst Biol. 56:17–24.

Kutschera V.E., Bidon T., Hailer F., Rodi J.L., Fain S.R., Janke A. 2014. Bears in a forest of gene trees: phylogenetic inference is complicated by incomplete lineage sorting and gene flow. Mol Biol Evol. 31:2004–2017.

Lande R. 1979. Quantitative genetic analysis of multivariate evolution, applied to brain: body size allometry. Evolution. :402–416.

Leaché A.D., Harris R.B., Rannala B., Yang Z. 2014. The influence of gene flow on species tree estimation: a simulation study. Syst Biol. 63:17–30.

Lemos B., Bettencourt B.R., Meiklejohn C.D., Hartl D.L.. 2005. Evolution of proteins and gene expression levels are coupled in Drosophila and are independently associated with mRNA abundance, protein length, and number of protein-protein interactions. Mol. Biol. Evol. 22:1345– 1354.

Liu L., Xi Z., Wu S., Davis C.C., Edwards S. V. 2015. Estimating phylogenetic trees from genome-scale data. Ann N Y Acad Sci. 1360:36–53.

Liu L., Yu L. 2010. Phybase: an R package for species tree analysis. Bioinformatics. 26:962–963.

Long C., Kubatko L. 2018. The effect of gene flow on coalescent-based species-tree inference. Syst Biol. 67:770–785.

Maddison W.P. 1997. Gene trees in species trees. Syst Biol. 46:523–536.

Maddison W.P., FitzJohn R.G. 2015a. The unsolved challenge to phylogenetic correlation tests for categorical characters. Syst Biol. 64:127–136.

Martins E.P. 1996. Phylogenies, spatial autoregression, and the comparative method: a computer simulation test. Evolution. 50:1750–1765.

Martins E.P., Garland Jr T. 1991. Phylogenetic analyses of the correlated evolution of continuous characters: a simulation study. Evolution. 45:534–557.

Martins E.P., Hansen T.F. 1997. Phylogenies and the comparative method: a general approach to incorporating phylogenetic information into the analysis of interspecific data. Am Nat. 149:646– 667.

Mazel F., Davies T.J., Georges D., Lavergne S., Thuiller W., Peres-Neto P.R. 2016. Improving phylogenetic regression under complex evolutionary models. Ecology. 97:286–293.

Mendes F.K., Fuentes-González J.A., Schraiber J.G., Hahn M.W. 2018. A multispecies coalescent model for quantitative traits. Elife. 7:e36482.

Mendes F.K., Hahn M.W. 2016. Gene tree discordance causes apparent substitution rate variation. Syst Biol. 65:711–721.

Mendes F.K., Hahn Y., Hahn M.W. 2016. Gene tree discordance can generate patterns of diminishing convergence over time. Mol Biol Evol. 33:3299–3307.

Mendes F.K., Livera A.P., Hahn M.W. 2019. The perils of intralocus recombination for inferences of molecular convergence. Philosophical Transactions of the Royal Society B. 374:20180244.

Mount G.G., Brown J.M. 2022. Comparing likelihood ratios to understand genome-wide variation in phylogenetic support. Syst Biol. 71:973–985.

Moyers Arévalo R.L., Amador L.I., Almeida F.C., Giannini, N.P. 2020. Evolution of body mass in bats: insights from a large supermatrix phylogeny. Journal of Mammalian Evolution. 27:123–138.

Navarro Gonzalez J., Zweig A.S., Speir M.L., Schmelter D., Rosenbloom K.R., et al. 2021. The UCSC Genome Browser database: 2021 update. Nucleic Acids Res. 49:D1046–D1057.

Nichols R. 2001. Gene trees and species trees are not the same. Trends Ecol Evol. 16:358–364.

O’Meara B.C., Ané C., Sanderson M.J., Wainwright P.C. 2006. Testing for different rates of continuous trait evolution using likelihood. Evolution (N Y). 60:922–933.

Pagel M. 1997. Inferring evolutionary processes from phylogenies. Zool Scr. 26:331–348.

Pagel M. 1999b. Inferring the historical patterns of biological evolution. Nature. 401:877–884.

Pál C., Papp B., Hurst L.D. 2001. Highly expressed genes in yeast evolve slowly. Genetics. 158:927– 931.

Pardo L. 2005. Statistical inference based on divergence measures. Boca Raton, FL: Chapman and Hall/CRC.

Paradis E., Schliep K. 2019. Ape 5.0: an environment for modern phylogenetics and evolutionary analyses in R. Bioinformatics. 35:526–528.

Pease J.B., Haak D.C., Hahn M.W., Moyle L.C. 2016. Phylogenomics reveals three sources of adaptive variation during a rapid radiation. PLoS Biol. 14:e1002379.

Pei J., Wu Y. 2017. STELLS2: fast and accurate coalescent-based maximum likelihood inference of species trees from gene tree topologies. Bioinformatics. 33:1789–1797.

Pennell M.W., Eastman J.M., Slater G.J., Brown J.W., Uyeda J.C., FitzJohn R.G., Alfaro M.E., Harmon L.J. 2014a. geiger v2. 0: an expanded suite of methods for fitting macroevolutionary models to phylogenetic trees. Bioinformatics. 30:2216–2218.

Pennell M.W., Harmon L.J. 2013. An integrative view of phylogenetic comparative methods: connections to population genetics, community ecology, and paleobiology. Ann N Y Acad Sci. 1289:90–105.

Pinheiro J., Bates D., DebRoy S., Sarkar, D., Heisterkamp S., Van Willigen B., Maintainer R., 2017. Package ‘nlme’. Linear and nonlinear mixed effects models. 3: 274.

Pollard D.A., Iyer V.N., Moses A.M., Eisen M.B. 2006. Widespread discordance of gene trees with species tree in Drosophila: evidence for incomplete lineage sorting. PLoS Genet. 2:e173.

Revell L.J. 2010. Phylogenetic signal and linear regression on species data. Methods in Ecology and Evolution. 1:319–329.

Revell L.J. 2012. phytools: an R package for phylogenetic comparative biology (and other things). Methods Ecol Evol. 3:217–223.

Revell L.J., Harmon L.J., Collar D.C. 2008. Phylogenetic signal, evolutionary process, and rate. Syst Biol. 57:591–601.

Robinson D.F., Foulds, L.R. 1981. Comparison of phylogenetic trees. Math. Biosci., 53:131–147.

Robinson M.L., Hahn P.G., Inouye B.D., Underwood N., Whitehead S.R., Abbott K.C., Bruna E.M., Cacho N.I., Dyer L.A., Abdala-Roberts L., Herbivory Variability Network, 2023. Plant size, latitude, and phylogeny explain within-population variability in herbivory. Science, 382:679–683.

Rousseeuw P., Yohai V. 1984. Robust regression by means of S-estimators. Robust and nonlinear time series analysis. New York. Springer. p. 256–272.

Rohlf F.J. 2001a. Comparative methods for the analysis of continuous variables: Geometric interpretations. Evolution (N Y). 55:2143–2160.

Ross C.F., Henneberg M., Ravosa M.J., Richard, S., 2004. Curvilinear, geometric and phylogenetic modeling of basicranial flexion: is it adaptive, is it constrained? Journal of Human Evolution. 46:185–213.

Sanford G.M., Lutterschmidt W.I., Hutchison V.H. 2002a. The comparative method revisited. Bioscience. 52:830–836.

Schluter D. 1995. Uncertainty in ancient phylogenies. Nature. 377:108–109.

Shen X.-X., Hittinger C.T., Rokas A. 2017. Contentious relationships in phylogenomic studies can be driven by a handful of genes. Nat Ecol Evol. 1:126.

Shriver, M.D., Kennedy, G.C., Parra, E.J., Lawson, H.A., Sonpar, V., Huang, J., Akey, J.M. Jones, K.W. 2004. The genomic distribution of population substructure in four populations using 8,525 autosomal SNPs. Hum. Genomics. 1:274–286.

Slatkin M., Pollack J.L. 2008. Subdivision in an ancestral species creates asymmetry in gene trees. Mol Biol Evol. 25:2241–2246.

Solís-Lemus C., Yang M., Ané C. 2016. Inconsistency of species tree methods under gene flow. Syst Biol. 65:843–851.

Stadler T. 2011. Simulating trees with a fixed number of extant species. Systematic biology. 60: 676– 684.

Stone E.A. 2011. Why the phylogenetic regression appears robust to tree misspecification. Syst Biol. 60:245–260.

Subramanian S, Kumar S. 2004. Gene expression intensity shapes evolutionary rates of the proteins encoded by the vertebrate genome. Genetics. 168:373–381

Symonds M.R.E. 2002a. The effects of topological inaccuracy in evolutionary trees on the phylogenetic comparative method of independent contrasts. Syst Biol. 51:541–553.

Symonds M.R.E., Blomberg S.P. 2014. A primer on phylogenetic generalised least squares. Modern phylogenetic comparative methods and their application in evolutionary biology. Springer. 105– 130.

Tian Y., Kubatko L.S. 2016. Distribution of coalescent histories under the coalescent model with gene flow. Mol Phylogenet Evol. 105:177–192.

Uyeda J.C., Zenil-Ferguson R., Pennell M.W. 2018a. Rethinking phylogenetic comparative methods. Syst Biol. 67:1091–1109.

Villemereuil P. de, Wells J.A., Edwards R.D., Blomberg S.P. 2012. Bayesian models for comparative analysis integrating phylogenetic uncertainty. BMC Evol Biol. 12:1–16.

Walker J.F., Brown J.W., Smith S.A. 2018. Analyzing contentious relationships and outlier genes in phylogenomics. Syst Biol. 67:916–924.

Wascher M., Kubatko L.S. 2023. On the effects of selection and mutation on species tree inference. Mol Phylogenet Evol. 179:107650.

Weaver L.N., Wilson G.P. 2021. Shape disparity in the blade-like premolars of multituberculate mammals: functional constraints and the evolution of herbivory. Journal of Mammalogy. 102: 967–985.

Yi X., Liang Y., Huerta-Sanchez E., Jin X., Cuo Z.X.P., Pool J.E., Xu X., Jiang H., Vinckenbosch N., Korneliussen T.S., Zheng, H. 2010. Sequencing of 50 human exomes reveals adaptation to high altitude. Science. 329:75–78.

Yu Y., Than C., Degnan J.H., Nakhleh L. 2011. Coalescent histories on phylogenetic networks and detection of hybridization despite incomplete lineage sorting. Syst Biol. 60:138–149.

Yule G.U. 1925a. A mathematical theory of evolution, based on the conclusions of Dr. JC Willis, FR S. Philosophical transactions of the Royal Society of London. Series B, containing papers of a biological character. 213:21–87.

Zhang R., Drummond A.J., Mendes F.K., 2021. Fast Bayesian inference of phylogenies from multiple continuous characters. bioRxiv. 2021–04.

Zhang G., Li C., Li Q., Li B., Larkin D.M., Lee C., Storz J.F., Antunes A., Greenwold M.J., Meredith R.W. 2014. Comparative genomics reveals insights into avian genome evolution and adaptation. Science (1979). 346:1311–1320.

