## Supplementary material for "A tale of too many trees: a conundrum for phylogenetic regression": Fig. S1

### SUPPLEMENTARY FIGURES

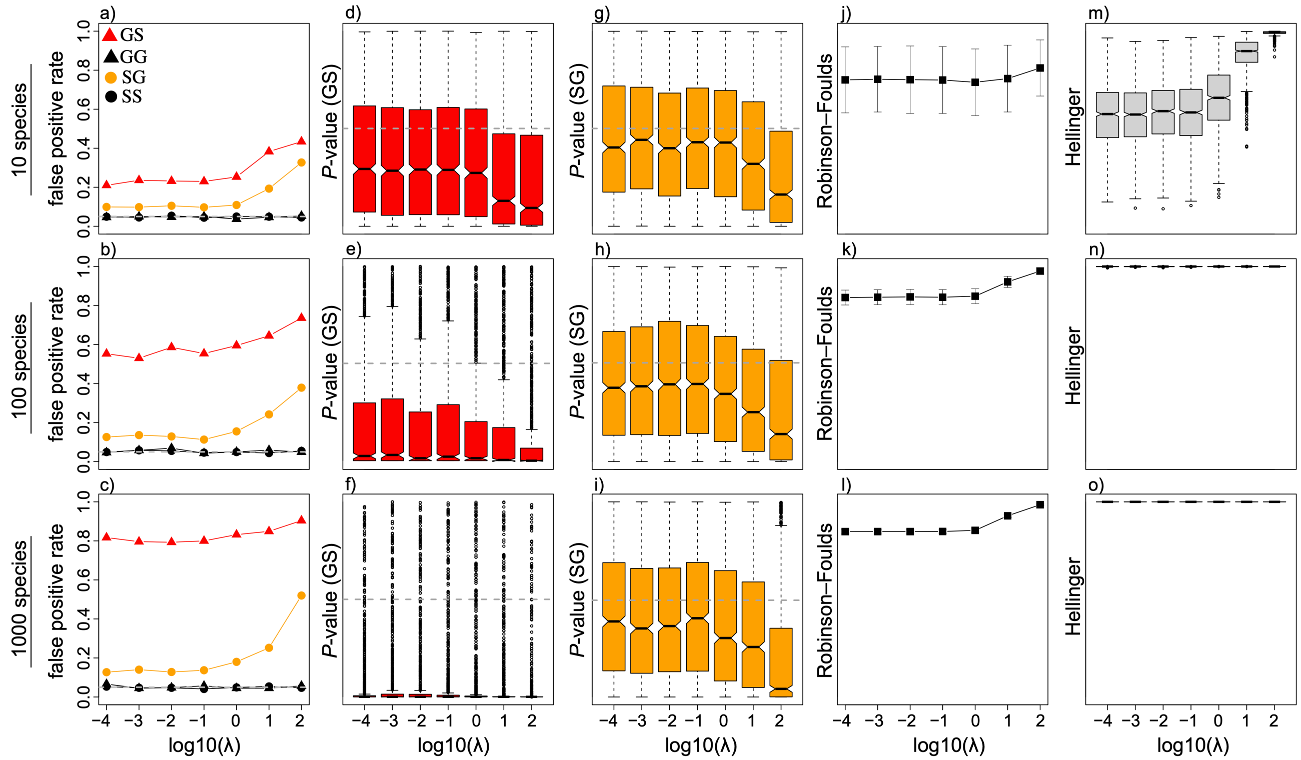


FIGURE S1. Impacts of mismatched phylogenetic regression. Estimates of the false positive rate (a-c), *P*-value distributions for GS (d-f), *P*-value distributions for SG (g-i), means and standard deviations of Robinson-Foulds topology distances (j-l), and probabilistic Hellinger distances (m-o) between gene trees and species trees from simulations including of 10 species (top row), 100 species (middle row), and 1000 species (bottom row) for birth-death simulations with birth rate $\lambda$, death rate $\lambda/2$, and root age of one coalescent unit. The two traits were statistically independent ($\beta = 0$) for all simulations. Dashed horizontal lines mark the commonly used false positive rate $\alpha = 0.05$ in panels a-c, median *P*-values taken from matched GG scenarios in panels d-f, and median *P*-values for matched SS scenarios in panels g-1. The *y*-axis ranges from zero to one in all panels.

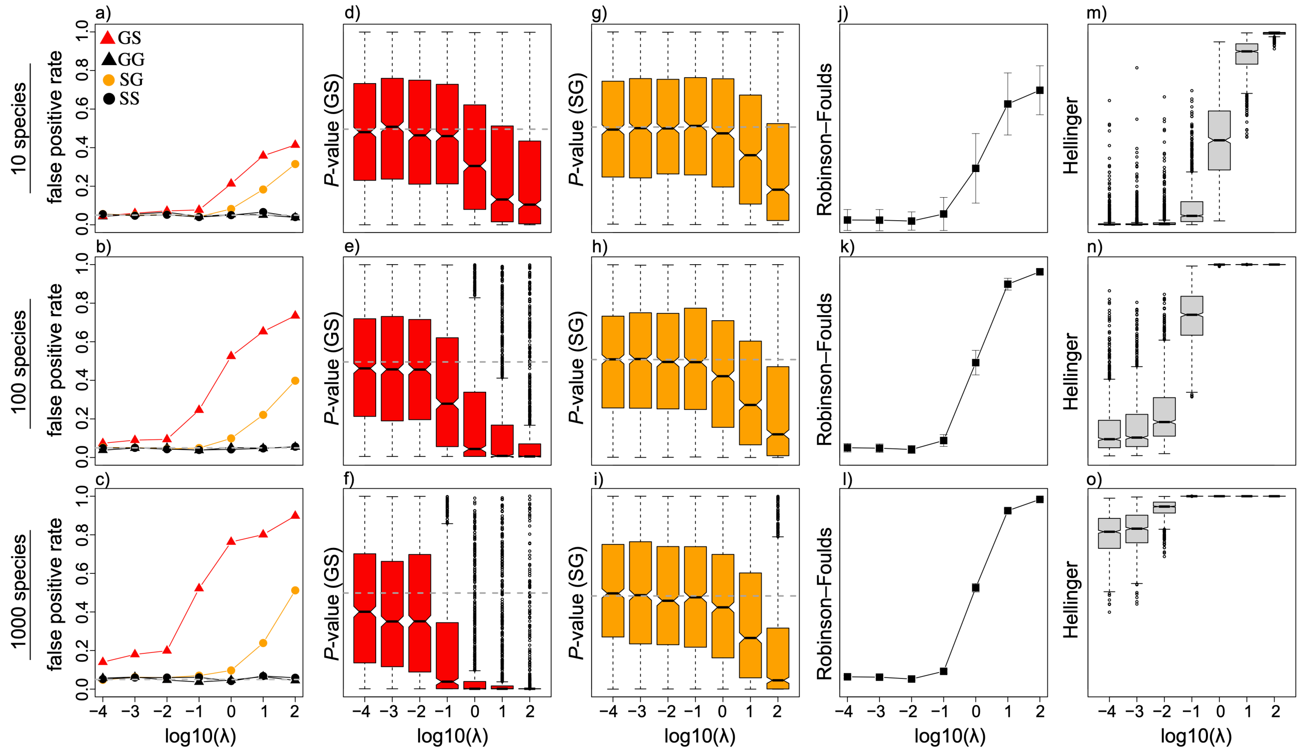


FIGURE S2. Impacts of mismatched phylogenetic regression. Estimates of the false positive rate (a-c), $P$-value distributions for GS (d-f), $P$-value distributions for SG (g-i), means and standard deviations of Robinson-Foulds topology distances (j-l), and probabilistic Hellinger distances (m-o) between gene trees and species trees from simulations including of 10 species (top row), 100 species (middle row), and 1000 species (bottom row) for birth-death simulations with birth rate $\lambda$, death rate $\lambda/2$, and root age of 100 coalescent units. The two traits were statistically independent ($\beta = 0$) for all simulations. Dashed horizontal lines mark the commonly used false positive rate $\alpha = 0.05$ in panels a-c, median $P$-values taken from matched GG scenarios in panels d-f, and median $P$-values for matched SS scenarios in panels g-1. The $y$-axis ranges from zero to one in all panels.

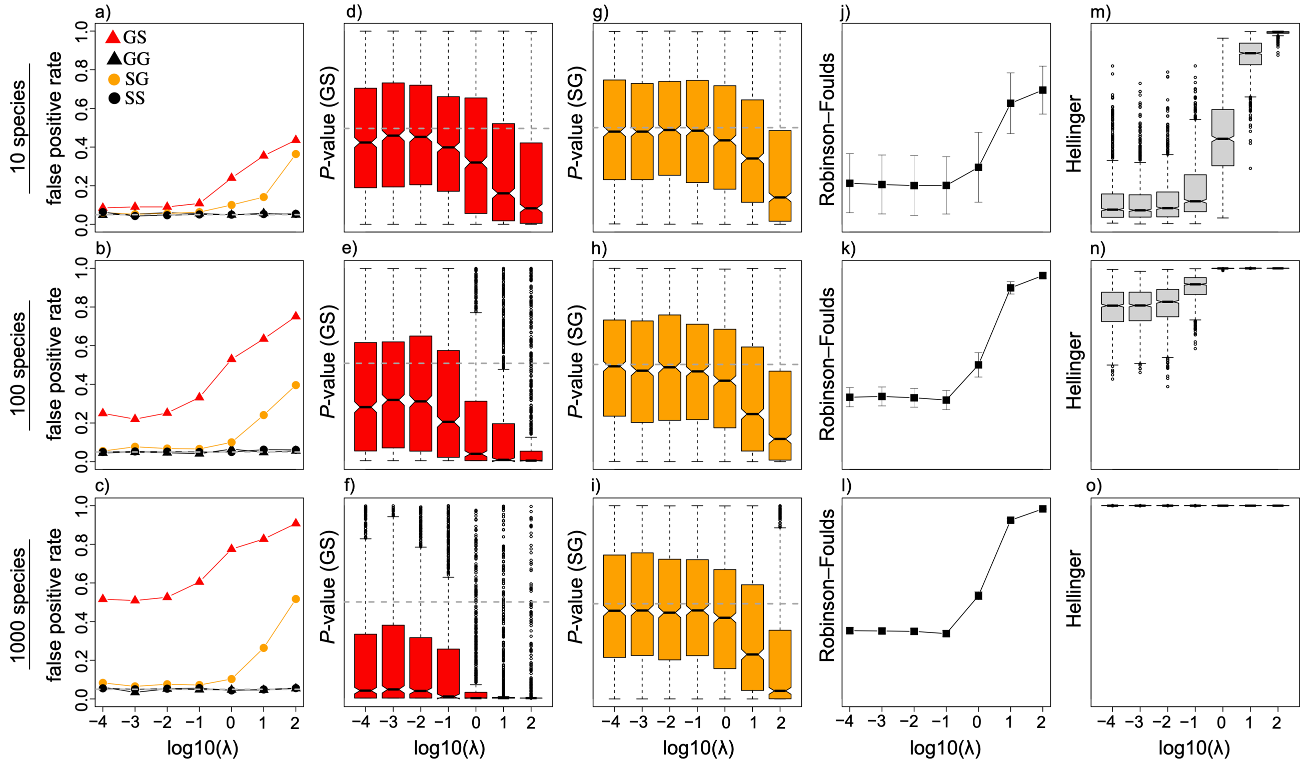


FIGURE S3. Impacts of mismatched phylogenetic regression. Estimates of the false positive rate (a-c), $P$-value distributions for GS (d-f), $P$-value distributions for SG (g-i), means and standard deviations of Robinson-Foulds topology distances (j-l), and probabilistic Hellinger distances (m-o) between gene trees and species trees from simulations including of 10 species (top row), 100 species (middle row), and 1000 species (bottom row) for pure-birth simulations with birth rate $\lambda$ and root age of 10 coalescent units. The two traits were statistically independent ($\beta = 0$) for all simulations. Dashed horizontal lines mark the commonly used false positive rate $\alpha = 0.05$ in panels a-c, median $P$-values taken from matched GG scenarios in panels d-f, and median $P$-values for matched SS scenarios in panels g-1. The $y$-axis ranges from zero to one in all panels.

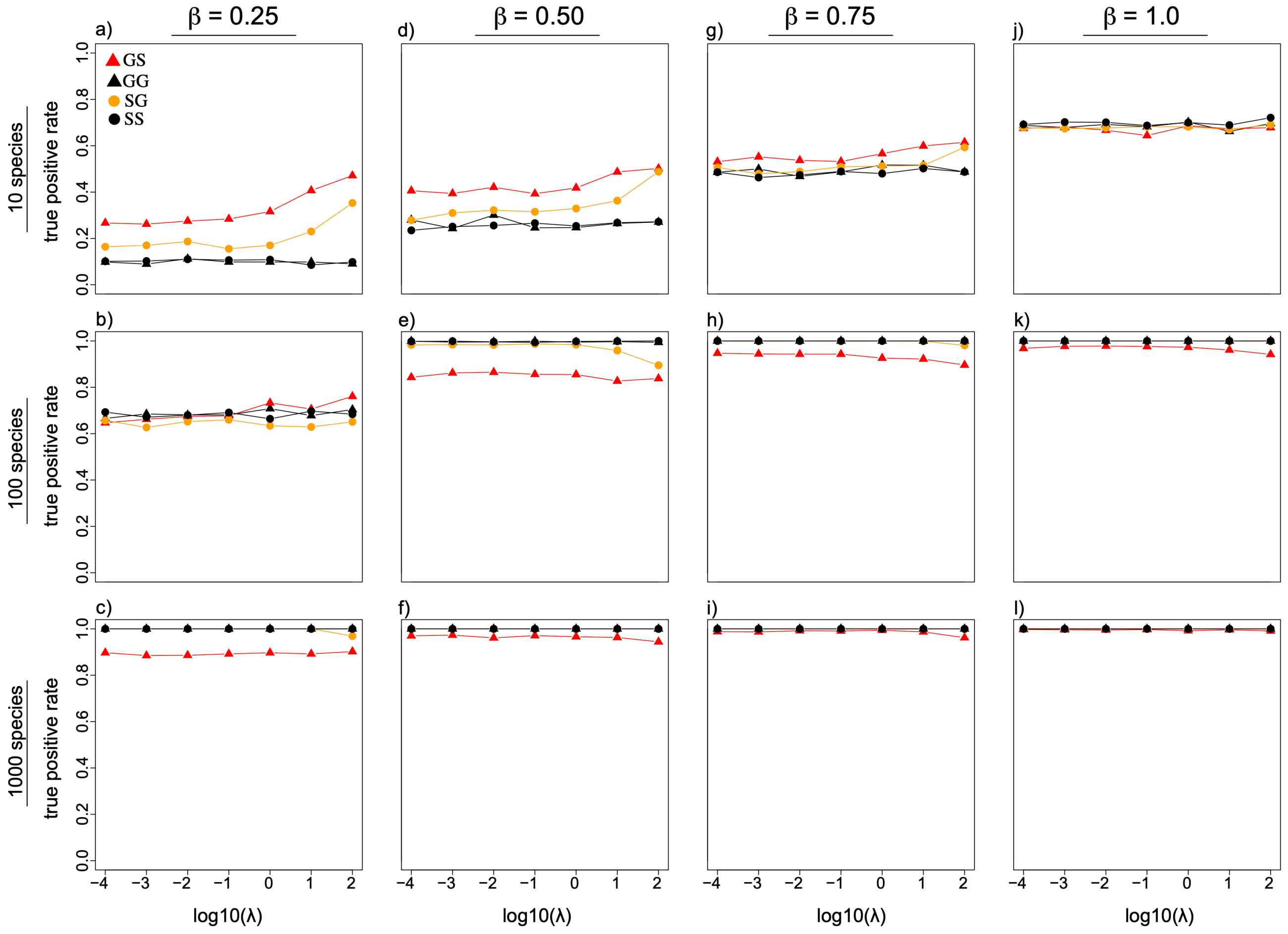


FIGURE S4. Impacts of mismatched phylogenetic regression on statistical power to detect trait associations. Estimates of the true positive rate for 10 species (top row), 100 species (middle row), and 1000 species (bottom row) for birth-death simulations with birth rate $\lambda$, death rate $\lambda/2$, and root age of one coalescent unit. Results shown for $\beta = 0.25$ (a-c), $\beta = 0.50$ (d-f), $\beta = 0.75$ (g-i), and $\beta = 1.0$ (j-l).

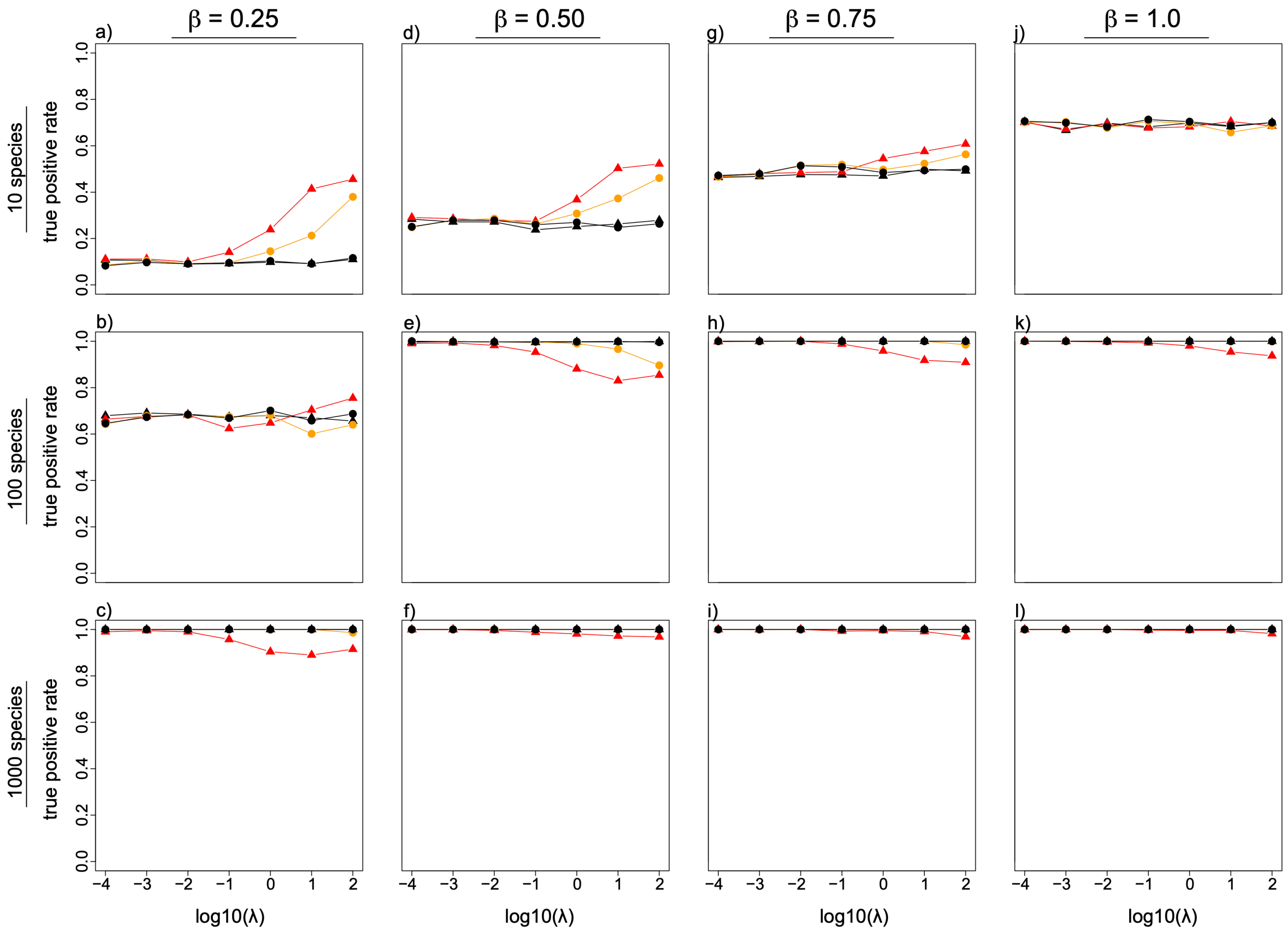


FIGURE S5. Impacts of mismatched phylogenetic regression on statistical power to detect trait associations. Estimates of the true positive rate for 10 species (top row), 100 species (middle row), and 1000 species (bottom row) for birth-death simulations with birth rate $\lambda$, death rate $\lambda/2$, and root age of 100 coalescent units. Results shown for $\beta = 0.25$ (a-c), $\beta = 0.50$ (d-f), $\beta = 0.75$ (g-i), and $\beta = 1.0$ (j-l).

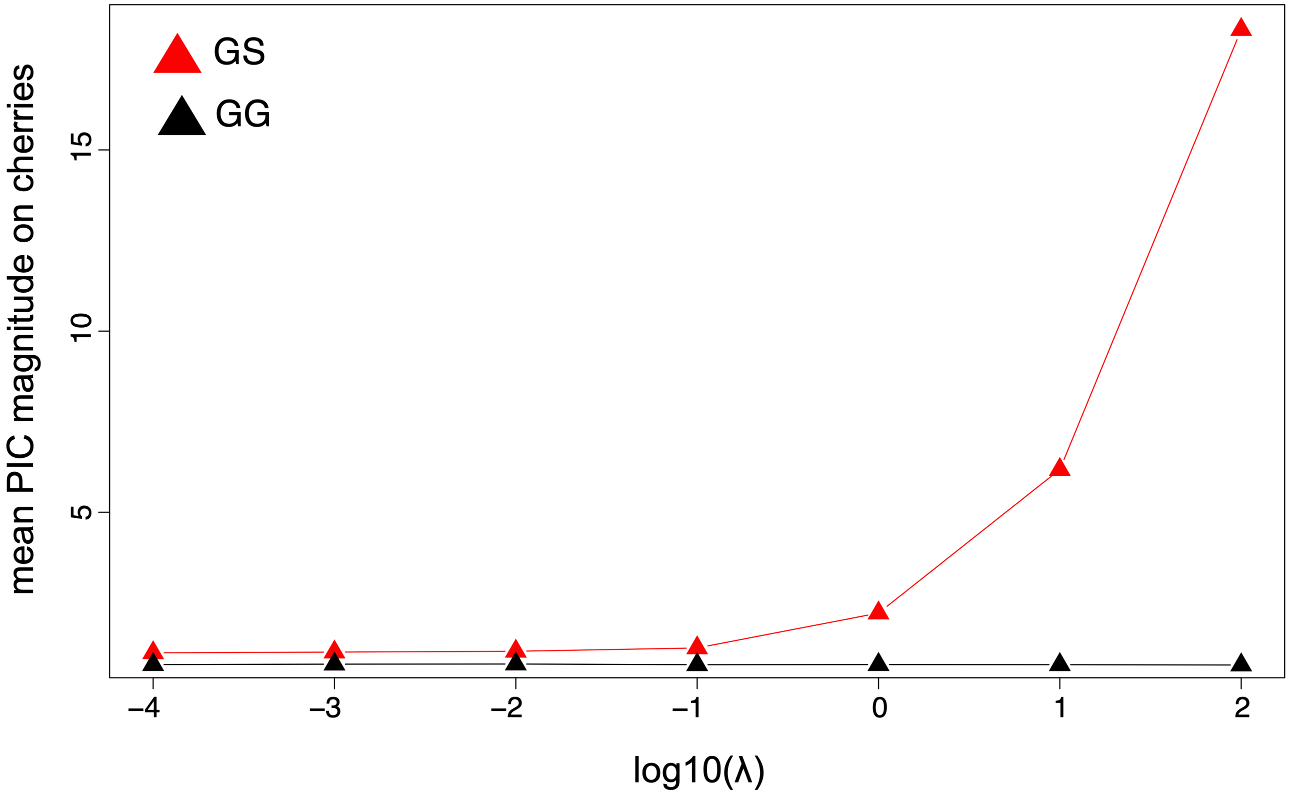


FIGURE S6. The relationship between mean magnitudes of PICs for cherries (i.e., clades with two extant tips) and speciation rates for GS (red lines) and GG (black lines) scenarios. Analyses shown for simulations including 10 species for birth-death simulations with birth rate $\lambda$, death rate $\lambda/2$, and root age of 10 coalescent units.
